# Simvastatin Therapy Attenuates Memory Deficits that Associate with Brain Monocyte Infiltration in Chronic Hypercholesterolemia

**DOI:** 10.1101/2020.05.15.098236

**Authors:** Nicholas Don-Doncow, Lotte Vanherle, Frank Matthes, Sine Kragh Petersen, Hana Matuskova, Sara Rattik, Anetta Härtlova, Anja Meissner

## Abstract

Evidence associates cardiovascular risk factors with unfavorable systemic and neuro-inflammation and cognitive decline in the elderly. Cardiovascular therapeutics (e.g., statins and anti-hypertensives) possess immune-modulatory functions in parallel to their cholesterol- or blood pressure (BP)-lowering properties. How their ability to modify immune responses affects cognitive function is unknown. Here, we examined the effect of chronic hypercholesterolemia on inflammation and memory function in Apolipoprotein E (ApoE) knockout mice and normocholesterolemic wild-type mice. Chronic hypercholesterolemia that was accompanied by moderate blood pressure elevations associated with apparent immune system activation characterized by increases in circulating pro-inflammatory Ly6Chi monocytes in ApoE^-/-^ mice. The persistent low-grade immune activation that is associated with chronic hypercholesterolemia facilitates the infiltration of pro-inflammatory Ly6Chi monocytes into the brain of aged ApoE^-/-^ but not wild-type mice, and links to memory dysfunction. Therapeutic cholesterol-lowering through simvastatin reduced systemic and neuro-inflammation, and the occurrence of memory deficits in aged ApoE^-/-^ mice with chronic hypercholesterolemia. BP-lowering therapy alone (i.e., hydralazine) attenuated some neuro-inflammatory signatures but not the occurrence of memory deficits. Our study suggests a link between chronic hypercholesterolemia, myeloid cell activation and neuro-inflammation with memory impairment and encourages cholesterol-lowering therapy as safe strategy to control hypercholesterolemia-associated memory decline during ageing.

## Introduction

Cognitive decline is an increasingly common problem that progresses with age. By gradually interfering with daily functioning and well-being, it poses an enormous burden on affected people and their environment ^1,2^. In recent years, the apparent role of cardiovascular risk factors as important modifiable elements in the development of cognitive decline has become the subject of clinical as well as pre-clinical research efforts ^2-4^. There exists a prominent link between mid-life chronic hypercholesterolemia and/or hypertension and the development of dementia later in life ^5,6^. As a consequence, the management of cardiovascular risk factors not only serves to prevent often detrimental acute consequences of hypertension or dyslipidemia, but also to minimize potential adverse cognitive outcomes later on. A number of observational and randomized studies reported beneficial effects of antihypertensive therapies on cognitive outcome ^7-9^. Nonetheless, existing distinctions regarding effectiveness between different classes of hypertension medication have been shown ^7^. Likewise, the use of statins in controlling risk factors for dementia prevention or treatment is controversially discussed. Their therapeutic efficacy that initially emerged from findings showing its positive effects on various factors related to memory function ^10,11^ was increasingly argued as case studies raised concerns regarding a contributory role of statins in development of cognitive problems ^12,13^. Due to the discrepancies observed between studies and the lack of knowledge regarding underlying mechanisms, strategies to combat cognitive impairment resulting from cardiovascular risk factors or cardiovascular disease (CVD) are yet to be established.

Immune system alterations and inflammation are not only hallmarks of many CVDs and their risk factors but have increasingly been recognized as contributors to impaired cognitive function in older people ^14-16^. Pharmaceuticals directed against inflammation evidently improve cognitive alterations associated with normal aging, CVDs, and classical neurodegenerative diseases ^17,18^. Multiple drugs commonly used in the cardiovascular field possess immune-modulatory actions in parallel to their principal cholesterol- or blood pressure (BP)-lowering effects ^19,20^. One of the bottlenecks hampering the targeted use of CVD therapeutics as an approach to minimize CVD-associated cognitive impairment is the lacking understanding of specific inflammatory signatures that relate to cognitive impairment in patients with increased cardiovascular burden.

The herein presented study investigated the effect of chronic exposure to elevated plasma cholesterol levels on systemic and neuro-inflammation, and memory function in ageing mice. Using mice deficient in Apolipoprotein E (ApoE), which normally facilitates cholesterol clearance in the liver and thus, controls plasma cholesterol levels, allowed us to mimic a chronically hypercholesterolemic state under normal dietary conditions. To study the effect of chronic hypercholesterolemia (i.e., higher-than-normal plasma cholesterol already apparent at 4-months of age) on systemic and neuro-inflammation and memory performance, we compared ApoE^-/-^ mice to normocholesterolemic wild-type (WT) mice. We tested the efficacy of CVD therapeutics as treatment for memory impairment associated with the increased cardiovascular burden in aged ApoE^-/-^ mice (i.e., increased plasma cholesterol and higher-than-normal BP). To this end, we therapeutically administered the statin simvastatin that has been attributed a strong anti-inflammatory character by regulating proliferation and activation of macrophages ^21,22^, and/or the smooth muscle relaxant agent hydralazine, known to reduce leukocyte migration in spontaneously hypertensive rats ^23^. Our findings provide evidence that early life exposure to chronically high cholesterol levels (i.e., apparent at 4-months of age) rather than increases in plasma cholesterol and BP later in life negatively affect memory performance. We underscore a role of myeloid cell activation in the development of neuro-inflammation associated with chronic hypercholesterolemia as well as cell type-specific drug effects that may limit their efficacy.

## Results

### Chronic hypercholesterolemia contributes to the development of memory impairment

To investigate whether chronic exposure to higher-than-normal plasma cholesterol levels affects important brain functions (i.e., cognition) during ageing, we tested memory function in normocholesterolemic WT and hypercholesterolemic ApoE^-/-^ mice at 4 and 12 months of age (**Fig.1a**). In accordance with the increased cardiovascular burden in aged ApoE^-/-^ compared to WT mice (i.e., chronic hypercholesterolemia as evident from higher-than-normal plasma cholesterol already apparent at 4 months of age and higher systolic BP at 12 months of age in ApoE^-/-^ mice; **Table 1**), long-term (hippocampus-dependent) memory function deteriorated with age in ApoE^-/-^ mice, resulting in significantly lower recognition indices (RI) compared to age-matched WT controls (RI≤0.5; **Fig.1b**). Although neither aged WT nor aged ApoE^-/-^ mice presented with signs of short-term (rhino-cortical) memory impairment when compared to their young controls (**Supplementary Table 1**), spatial short-term memory was compromised in aged ApoE^-/-^ mice as evident by a lower RI obtained in an object placement test (RI≤0.5; **Fig.1c**). When testing for age-related changes in pre- and post-synaptic proteins in whole brain lysates of ApoE^-/-^ mice, no differences in expression of synaptosomal associated protein 25 (SNAP-25) or postsynaptic density protein 95 (PSD-95) were observed (**Fig.1d**). In accordance with previous studies ^24^, however, ageing affected hippocampal but not cortical PSD-95 mRNA expression in ApoE^-/-^ mice (**Fig.1e**). Taken together, these data imply brain region-specific negative effects of early and chronic hypercholesterolemia on memory function later in life.

**Figure 1:**
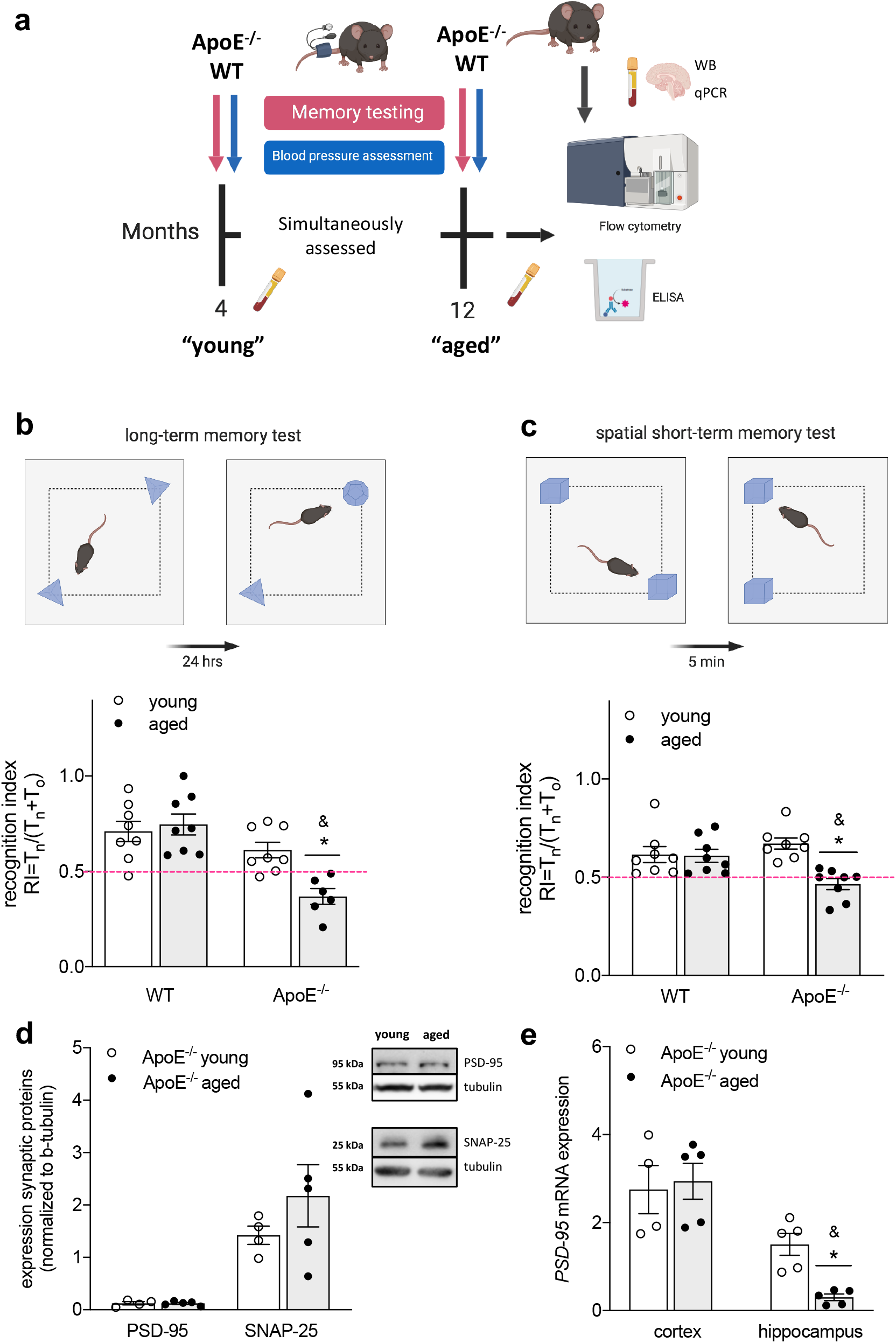
Chronic hypercholesterolemia impairs memory function. **(a)** BP and memory function were tested in 4- and 12-months old ApoE^-/-^ and WT mice simultaneously. At termination, immune status in blood and brain was assessed using flow cytometry, ELISA and qPCR. Synaptic marker expression was determined by WB and qPCR. This approach aimed at characterizing aging in the ApoE^-/-^ model in respect to plasma cholesterol levels, BP, systemic and neuro-inflammation as well as memory performance. **(b)** Schematic illustration of novel object recognition task with 24-hrs delay interval. RI describing hippocampus-dependent long-term memory function in young and aged WT and ApoE^-/-^ mice (N=10 per group; some animals were excluded due to total object exploration times below 20s). The pink dotted line indicates the point where the animal has no preference for either object. **(c)** Schematic illustration of an object placement test with a 5-min delay interval. RI describing spatial short-term memory function in young and aged WT and ApoE^-/-^ mice (N=10 per group; some animals were excluded due to total object exploration times below 20s). The pink dotted line indicates the point where the animal has no preference for either object. **(d)** Western blot analysis displaying unchanged pre- and post-synaptic protein (SNAP-25 and PSD-95) expression levels in whole brain tissue lysates of young and aged ApoE^-/-^ mice (N=4 for young and N=5 for aged ApoE^-/-^). Western blots are derived from the same experiment and were processed in parallel. **(e)** Brain region-specific mRNA expression analysis revealing reduced PSD-95 expression levels in hippocampus of aged ApoE^-/-^ mice (N=4 for young cortex and N=5 for young hippocampus and aged cortex and hippocampus). ApoE – apolipoprotein E, BP – blood pressure, RI – Recognition index, SNAP-25 – synaptosomal associated protein 25, PSD-95 – postsynaptic density protein 95, WB – western blotting, WT – wild-type. Data expressed as mean ± SEM. N denotes number of independent biological replicates. In (**b)-(e)**, * denotes P ≤ 0.05 relative to young control of same genotype, & denotes P ≤ 0.05 relative to WT aged after two-way ANOVA followed by Sidak *post hoc* testing. All schematics were created with BioRender.

**Table 1:** Hemodynamic data and plasma cholesterol levels of WT and ApoE^-/-^ mice. Systolic blood pressure and heart rate were measured in conscious mice using tail cuff plethysmography. Heart – body weight ratio calculated from animal weight (g) and respective heart weight (mg). Cholesterol levels were determined in plasma obtained from anaesthetized mice by ELISA. ApoE – apolipoprotein E, BP_sys_ – systolic blood pressure, bpm – beats per minute, BW – body weight, HR – heart rate, WT – wild-type. *Values are expressed as mean +/- SEM. N denotes number of independent biological replicates. N=10 per group; * denotes P ≤ 0*.*05 relative to respective young, & denotes P ≤ 0*.*05 relative to aged mice of respective other group after two-way ANOVA followed by Sidak post hoc test*.

| Physiological parameter | ApoE<sup>-/-</sup> young (4 months) | ApoE<sup>-/-</sup> aged (12 months) | WT young (4 months) | WT aged (12 months) |
| --- | --- | --- | --- | --- |
| BP<sub>sys</sub> (mmHg) | 104.8 ± 2.8 | 133.2 ± 4.4* | 99.4 ± 3.4 | 114.6 ± 1.7* |
| HR (bpm) | 489.1 ± 9.1 | 557.2 ± 11.0* | 458.9 ± 17.7 | 556.9 ± 11.3* |
| Cholesterol (mg/L) | 98.3 ± 2.4 | 181.7 ± 6.1* | 74.2 ± 0.8& | 99.8 ± 2.6*& |
| Heart – body weight ratio | 5.03 ± 0.19 | 5.91 ± 0.26* | 5.34 ± 0.13 | 5.48 ± 0.27 |

### Pro-inflammatory monocytes infiltrate the brain of aged ApoE^-/-^ but not WT mice

In chronically hypercholesterolemic ApoE^-/-^ mice, ageing associated with apparent immune system activation as evident by a higher number of circulating Ly6Chi monocytes (**Fig.2a**) and elevated plasma levels of interleukin (IL)12/23 (**Fig.2b**). The latter is often secreted by activated monocytes/macrophages promoting T helper (Th) 1 and Th17 priming of T-cells ^25^, which secrete IL17 amongst other cytokines and are considered main contributors to tissue inflammation ^26^. Elevated IL17 plasma levels in aged ApoE^-/-^ mice (**Fig.2c**) led us to test immune cell infiltration into brain tissue. FACS analyses revealed an increase of Ly6Chi pro-inflammatory monocytes (**Fig.2d**) and a higher percentage of overall T-cells (positive for the pan-T-cell marker CD3) in brain tissue of aged vs. young ApoE^-/-^ mice (**Fig.2e**). Transcripts of key chemokines that regulate migration and infiltration of monocytes and macrophages, such as monocyte chemoattractant protein-1 (MCP-1) and chemokine ligand-2 (CXCL2) were drastically elevated in whole brain tissue of the aged cohort compared to young control mice (**Table 2**). Moreover, an mRNA expression pattern typical of a pro-inflammatory phenotype was predominant in whole brain tissue of aged ApoE^-/-^ mice compared to young ApoE^-/-^ controls (**Table 2**). The degree of cytokine expression was brain region-dependent as evidenced by a markedly higher hippocampal than cortical IL12 expression in aged ApoE^-/-^ mice compared to young controls (**Table 2**). IL23 also showed such region-specific expression patterns (**Table 2**). Besides, Arginase-1 (Arg-1) as commonly used marker for an alternatively activated macrophage phenotype presented with region-specific expression patterns: while cortical Arg-1 mRNA levels remained unaffected by age, the basally higher hippocampal Arg-1 expression significantly reduced with age (**Table 2**). Together, these data point to an apparent age-related inflammation in the hippocampus and may explain the more pronounced hippocampal-dependent memory impairment in aged ApoE^-/-^ mice.

**Figure 2:**
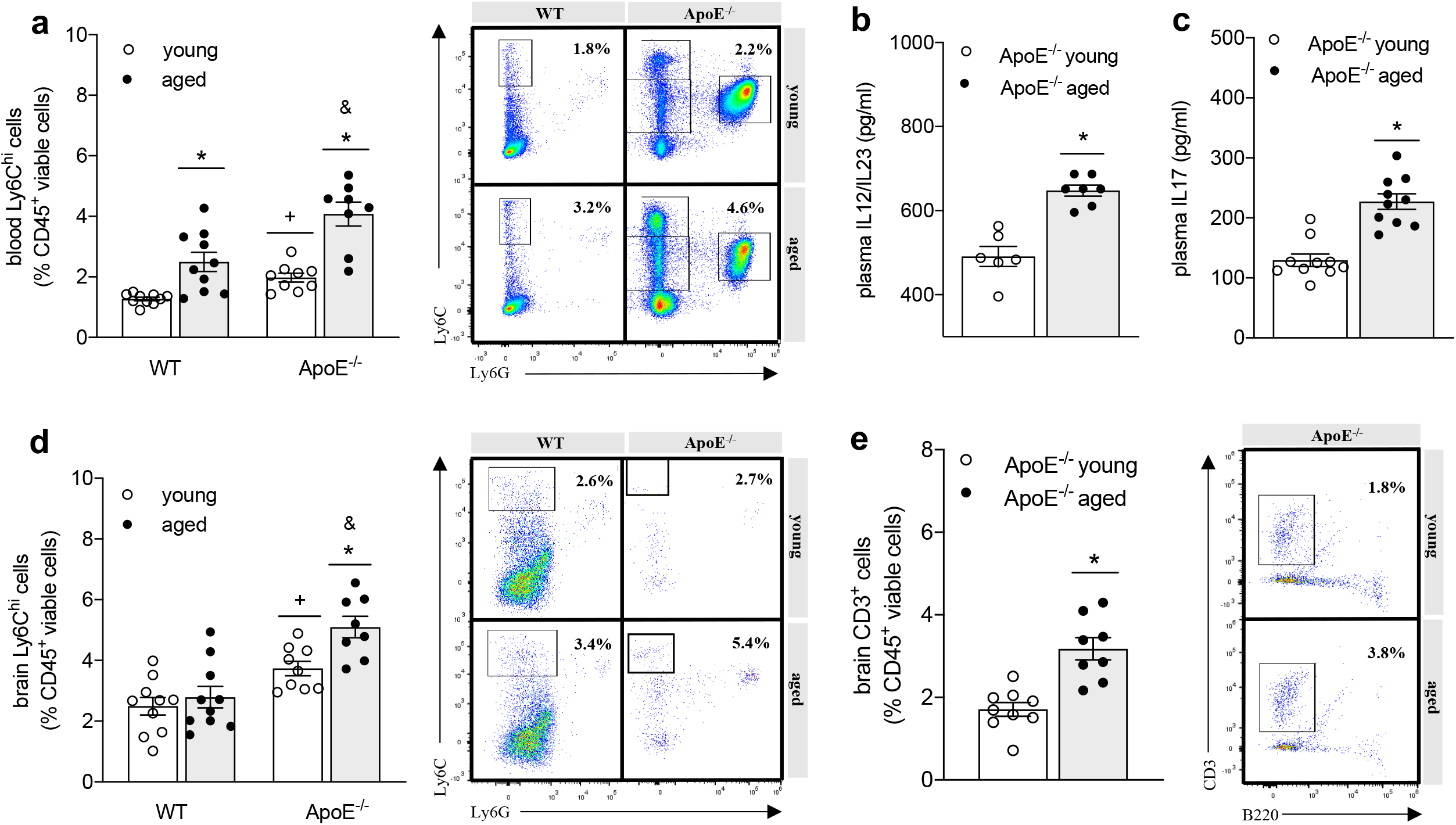
Immune system activation leads to leukocyte infiltration into the brain of aged ApoE^-/-^ mice. **(a)** Percentage of circulating Ly6Chi monocytes of young and aged WT (N=10 per group) compared to ApoE^-/-^ mice (N=8 and 9 per group) analyzed by flow cytometry and representative images showing dot plots for WT and ApoE^-/-^ mice. **(b)** Plasma levels of pro-inflammatory IL12/23 and **(c)** IL17 of young and aged ApoE^-/-^ mice determined by ELISA (N=10 per group; some samples failed to return a signal and were excluded from analysis). **(d)** Percentage of Ly6Chi monocytes in brain tissue of young and aged WT compared to ApoE^-/-^ mice analyzed by flow cytometry and representative images showing dot plots for WT and ApoE^-/-^ mice. (N=10 per group; some samples were excluded from analysis due to heavy myelin contamination). **(e)** Percentage of CD3+ T-cells in brain tissue of young and aged ApoE^-/-^ mice analyzed by flow cytometry and representative dot plots (N=10 per group; some samples were excluded from analysis due to heavy myelin contamination). ApoE – apolipoprotein E, IL - interleukin WT – wild-type. Data expressed as mean ± SEM. N denotes number of independent biological replicates. In **(a) & (d)**, * denotes P ≤ 0.05 relative to young control of same genotype, & denotes P ≤ 0.05 relative to WT aged, and + denotes P ≤ 0.05 relative to WT young after two-way ANOVA followed by Sidak *post hoc* testing; in **(b), (c) & (e)**,* denotes P ≤ 0.05 for single, unpaired comparisons.

**Table 2:** Brain mRNA expression of inflammatory markers in young and aged ApoE^-/-^ mice. mRNA expression of prominent inflammatory markers, including markers for classically and alternatively activated microglia were determined in whole brain tissue isolated from 4- and 12-months old ApoE^-/-^ mice using quantitative real-time PCR (6 top rows). In addition, selected markers were tested in cortex and hippocampus fractions. ApoE – apolipoprotein E, Arg-1 – arginase 1, CXCL – chemokine (C-X-C motif) ligand, IL – interleukin, iNOS – inducible nitric oxide synthase, MCP-1 – monocyte chemoattractant protein 1, TNF-α – tumor necrosis factor alpha. *Values are expressed as normalized to housekeeping gene expression and are shows as mean +/- SEM. N denotes number of independent biological replicates. N=5 mice per group; * denotes P ≤ 0*.*05 after single unpaired comparisons (two-tailed t-test)*.

| Target gene | ApoE<sup>-/-</sup> young<br>(4 months) | ApoE<sup>-/-</sup> aged<br>(12 months) |
| --- | --- | --- |
| whole brain |  |  |
| IL12 | 0.92 ± 0.07 | 5.12 ± 0.59* |
| MCP-1 | 0.56 ± 0.04 | 7.55 ± 0.68* |
| TNF-α | 1.04 ± 0.12 | 3.37 ± 0.23* |
| IL6 | 1.26 ± 0.17 | 3.29 ± 0.30* |
| CXCL2 | 0.68 ± 0.17 | 29.32 ± 7.97* |
| iNOS | 0.32 ± 0.03 | 3.81 ± 0.31* |
| cortex |  |  |
| IL12 | 1.02 ± 0.05 | 5.01 ± 1.22* |
| IL23 | 0.20 ± 0.07 | 0.40 ± 0.16 |
| Arg-1 | 0.52 ± 0.08 | 0.53 ± 0.09 |
| hippocampus |  |  |
| IL12 | 1.28 ± 0.56 | 11.84 ± 4.91* |
| IL23 | 0.51 ± 0.09 | 2.44 ± 0.89* |
| Arg-1 | 3.31 ± 0.66 | 1.64 ± 0.32* |

Although WT mice presented with an age-related increase of plasma cholesterol, BP (**Table 1**) and circulating Ly6Chi pro-inflammatory monocytes (**Fig.2a**), the percentage of Ly6Chi monocytes remained unaltered in the brain of aged WT mice (**Fig.2d**). Notably compared to WT mice, ApoE^-/-^ mice already showed a significantly higher proportion of circulating Ly6Chi cells at young age (**Fig.2a**), suggesting a persistent pro-inflammatory state in chronically hypercholesterolemic ApoE^-/-^ mice, which may promote immune infiltration into the brain with implications for memory function. Correspondingly, the proportion of brain Ly6Chi monocytes was higher in ApoE^-/-^ mice compared to age-matched WT mice (**Fig.2d**). Furthermore, brain tissue of aged ApoE^-/-^ mice presented with significantly higher transcripts levels of chemokines MCP-1 and CXCL2 compared to aged-matched WT controls (**Supplementary Table 2**). Taken together, these data suggest a key role for Ly6Chi monocyte activation and brain infiltration during chronic hypercholesterolemia where memory deficits were also observed.

### Cholesterol- and/or BP-lowering therapy attenuate neuro-inflammation in aged ApoE^-/-^ mice

As chronically hypercholesterolemic ApoE^-/-^ mice at 12 months of age presented with marginally elevated BP compared to 4 months old ApoE^-/-^ mice, we subjected aged ApoE^-/-^ mice (i.e., 12 months of age) to lipid-lowering or BP-lowering therapy for 8 consecutive weeks, using the statin simvastatin (0.5 mg/kg/d) or the smooth muscle relaxant hydralazine (5 mg/kg/d). Because we did not anticipate statin-mediated effects on BP, we also administered a combination of the two drugs to a group of animals (**Fig.3a**). The doses administered correspond to therapeutic doses used in the clinic (20 mg/kg/d and 10 mg/kg/d, respectively). All treatment strategies proved similarly effective in lowering the age-related elevated systolic BP: longitudinal BP measurements revealed markedly lower BP values in mice post treatment compared to the recorded pre-treatment values (**Supplementary Fig.1**). As expected, simvastatin treatment revealed significant cholesterol-lowering effects (**Table 3**). In line with other investigations ^27,28,^ simvastatin treatment markedly reduced the age-related elevated heart-to-body weight ratio in our model without significantly affecting heart rate (**Table 3**). Interestingly, simvastatin’s effects on cardiac hypertrophy and cholesterol were diminished in the presence of hydralazine (**Table 3**). Simvastatin revealed considerable anti-inflammatory capacity as evidenced by a reduction of circulating plasma IL12/23 levels (**Fig.3b**) and IL17 levels *in vivo* (**Fig.3c**). In the brain, simvastatin-treated mice revealed lower transcript levels of pro-inflammatory cytokines and chemokines, including IL6 and IL12, MCP-1 and CXCL2 (**Table 4**). Notably, hydralazine attenuated simvastatin’s effects for some of the cytokines when administered in combination (**Table 4**). Treatment affected brain regions to different degrees as supported by statistically significant differences in hippocampal but not cortical IL12 expression between simvastatin- and hydralazine-treated groups (**Table 4**). Interestingly, IL23 revealed a different expression profile with significantly lower cortical IL23 expression in simvastatin-containing treatment groups while hippocampal IL23 expression only significantly changed when mice received simvastatin alone (**Table 4**).

**Figure 3:**
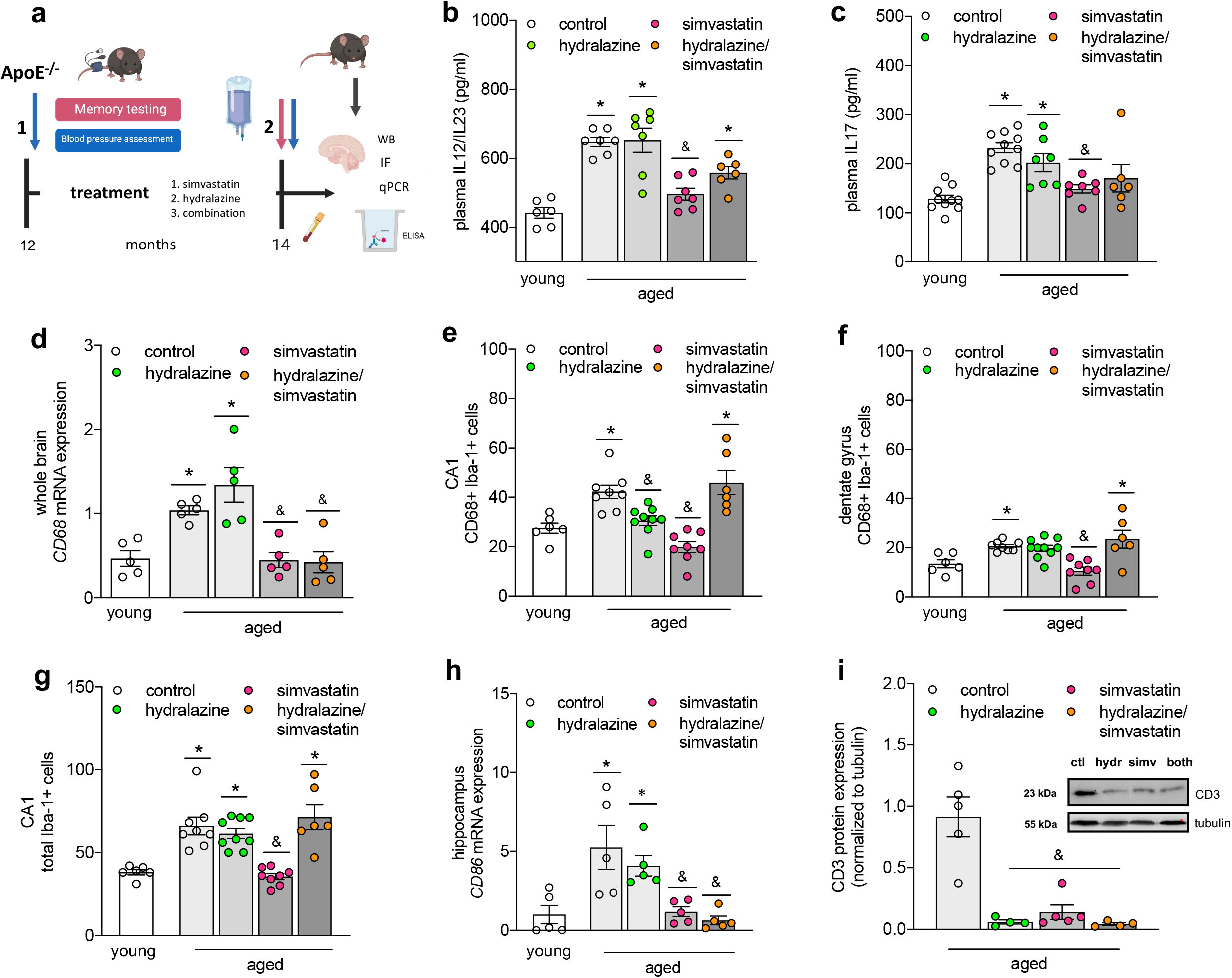
Simvastatin treatment attenuates systemic and neuro-inflammation in aged ApoE^-/-^ mice. **(a)** 12- months old ApoE^-/-^ mice (associated with chronic hypercholesterolemia, increased BP, augmented systemic and neuro-inflammation and memory deficits) were subjected to 8-weeks cholesterol-lowering therapy (i.e., simvastatin), BP-lowering therapy (i.e., hydralazine) or a combination of both. BP was assessed longitudinally to monitor the effect of the different treatment regimens. Memory function was assessed at 14 months of age. At termination, plasma cholesterol levels were determined by ELISA and systemic and neuro-inflammation were assessed by ELISA, qPCR, WB and immunofluorescence to characterize the effects of the different treatment regimens on the different signatures we identified to be altered in this model (i.e., plasma cholesterol, BP, inflammatory status and memory function). Schematic created with BioRender. **(b)** Effect of cholesterol- and BP-lowering treatment (hydralazine – green circle, simvastatin – pink circles, combination – orange circles) on plasma levels of pro-inflammatory IL12/23 and **(c)** IL17 determined by ELISA (N=10 per group and N=8 for aged + combination treatment; some samples failed to return a signal and were excluded from analysis). **(d)** Effect of cholesterol- and BP-lowering treatment on CD68 mRNA expression in whole brain extracts of aged ApoE^-/-^ mice (N=5 per group). **(e)** Effect of cholesterol- and BP-lowering treatment on the number of CD68+ Iba-1+ cells in the CA1 region of the hippocampus of aged ApoE^-/-^ mice (N=3 for young and aged + combination treatment; N=4 for aged control and simvastatin treatment; N=5 for hydralazine treatment; the two hemispheres were counted separately. **(f)** Effect of cholesterol- and BP-lowering treatment on the CD68+ Iba-1+ cells in the DG region of the hippocampus of aged ApoE^-/-^ mice (N=3 for young and aged + combination treatment; N=4 for aged control and simvastatin treatment; N=5 for hydralazine treatment; the two hemispheres were counted separately). **(g)** Effect of cholesterol- and BP-lowering treatment on the total Iba-1+ cell count in the CA1 region of the hippocampus of aged ApoE^-/-^ mice (N=3 for young and aged + combination treatment; N=4 for aged control and simvastatin treatment; N=5 for hydralazine treatment; the two hemispheres were counted separately except for a sample in the aged + hydralazine group where one hippocampus region was not analyzable). **(h)** Effect of cholesterol- and BP-lowering treatment on CD86 mRNA expression in the hippocampus of aged ApoE^-/-^ mice (N=5 per group). **(i)** Effect of cholesterol- and BP-lowering treatment on CD3 protein expression in whole brain lysates of aged ApoE^-/-^ mice (N=4 for hydralazine- and combination-treated groups and N=5 for control and simvastatin-treated groups). CD3 was not detected in brain tissue of young ApoE mice. Inset showing representative protein expression pattern. ApoE – apolipoprotein E, BP – blood pressure, CA – cornu ammonis, DG – dentate gyrus, Iba-1 – ionized calcium-binding adapter molecule-1, IL – interleukin, WB – western blotting. Data expressed as mean ± SEM. N denotes number of independent biological replicates. In (**b)-(d)** and **(f)-(i)**, * denotes P ≤ 0.05 relative to young control, & denotes P ≤ 0.05 relative to aged control after one-way ANOVA followed by Tukey’s *post hoc* testing. In **(e)**, * denotes P ≤ 0.05 relative to young control, & denotes P ≤ 0.05 relative to aged control after Kruskal-Wallis followed by Dunn’s *post hoc* testing.

**Table 3:** Hemodynamic data and plasma cholesterol levels of ApoE^-/-^ mice post treatment. Systolic blood pressure and heart rate were measured in conscious mice using tail cuff plethysmography. Heart – body weight ratio calculated from animal weight (g) and respective heart weight (mg). Cholesterol levels were determined in plasma obtained from anaesthetized mice. ApoE – apolipoprotein E, BP_sys_ – systolic blood pressure, bpm – beats per minute, BW – body weight, HR – heart rate. *Values are expressed as mean +/- SEM. N denotes number of independent biological replicates; N=10 per group and N=8 for aged ApoE*^*-/-*^ *treated with hydralazine/simvastatin combined; * denotes P ≤ 0*.*05 relative to young control, & denotes P ≤ 0*.*05 relative to aged group after one-way ANOVA followed by Tukey’s post hoc testing*.

| Physiological parameter | ApoE<sup>-/-</sup> young (4 months) | ApoE<sup>-/-</sup> aged (14 months) | ApoE<sup>-/-</sup> aged + hydralazine | ApoE<sup>-/-</sup> aged + simvastatin | ApoE<sup>-/-</sup> aged + hydralazine/simvastatin |
| --- | --- | --- | --- | --- | --- |
| BP<sub>sys</sub> (mmHg) | 107.3 ± 3.1 | 136.9 ± 5.8* | 110.1 ± 1.9& | 105.4 ± 1.6& | 112.6 ± 1.5& |
| HR (bpm) | 491.1 ± 9.7 | 569.9 ± 8.6* | 514.3 ± 10.9 | 512.2 ± 18.3 | 520.0 ± 12.9 |
| Cholesterol (mg/L) | 99.8 ± 4.3 | 184.7 ± 5.3* | 227.9 ± 13.6* | 108.7 ± 2.6*& | 187.4 ± 13.4* |
| Heart - body weight ratio | 4.59 ± 0.06 | 5.53 ± 0.21* | 5.27 ± 0.16* | 4.67 ± 0.56& | 5.31 ± 0.22* |

**Table 4:** Brain mRNA expression of pro-inflammatory cytokines and chemokines in ApoE^-/-^ mice after BP- and/or cholesterol-lowering treatment. mRNA expression of prominent inflammatory markers, including markers for classically and alternatively activated microglia were determined in whole brain tissue isolated from the different experimental groups using quantitative real-time PCR (6 top rows). In addition, selected markers were tested in cortex and hippocampus fractions. ApoE – apolipoprotein E, Arg-1 – arginase 1, BP – blood pressure, CXCL – chemokine (C-X-C motif) ligand, IL – interleukin, iNOS – inducible nitric oxide synthase, MCP-1 – monocyte chemoattractant protein 1, TNF-α – tumor necrosis factor alpha. *Values are expressed as normalized to housekeeping gene expression and shown as mean +/- SEM. N denotes number of independent biological replicates. N=5 mice per group; * denotes P ≤ 0*.*05 relative to young control, & denotes P ≤ 0*.*05 relative to aged control after one-way ANOVA followed by Tukey’s post hoc test*.

| target | ApoE<sup>-/-</sup> young<br>(4 months) | ApoE<sup>-/-</sup> aged (14<br>months) | ApoE<sup>-/-</sup> aged +<br>hydralazine | ApoE<sup>-/-</sup> aged +<br>simvastatin | ApoE<sup>-/-</sup> aged +<br>hydralazine/simvastatin |
| --- | --- | --- | --- | --- | --- |
| whole brain |  |  |  |  |  |
| IL12 | 0.98 ± 0.18 | 16.84 ± 2.29* | 5.74 ± 2.04*& | 1.39 ± 0.39<sup>&</sup> | 6.15 ± 2.48* |
| MCP-1 | 0.93 ± 0.19 | 9.11 ± 2.64* | 1.02 ± 0.13<sup>&</sup> | 1.32 ± 0.27<sup>&</sup> | 2.75 ± 0.53*& |
| TNF-α | 0.94 ± 0.09 | 7.33 ± 1.57* | 2.16 ± 0.38*& | 1.05 ± 0.28<sup>&</sup> | 2.19 ± 0.14*& |
| IL6 | 1.14 ± 0.13 | 2.95 ± 0.15* | 2.130± 0.37* | 1.62 ± 0.15<sup>&</sup> | 2.17 ± 0.23* |
| CXCL2 | 0.68 ± 0.17 | 29.32 ± 7.97* | 11.60 ± 7.85*& | 3.09 ± 2.15<sup>&</sup> | 31.55 ± 6.70* |
| iNOS | 1.00 ± 0.17 | 3.06 ± 0.79* | 2.38± 0.58*& | 1.89 ± 0.48<sup>&</sup> | 2.59 ± 0.62* |
| cortex |  |  |  |  |  |
| IL12 | 1.02 ± 0.05 | 5.07 ± 1.15* | 1.57 ± 0.54<sup>&</sup> | 0.97 ± 0.40<sup>&</sup> | 0.99 ± 0.19<sup>&</sup> |
| IL23 | 0.19 ± 0.11 | 1.16 ± 0.30* | 0.44 ± 0.16<sup>&</sup> | 0.09 ± 0.02<sup>&</sup> | 0.07 ± 0.03<sup>&</sup> |
| Arg-1 | 0.38 ± 0.23 | 0.11 ± 0.06 | 0.51 ± 0.26 | 0.43 ± 0.14 | 0.21 ± 0.14 |
| hippocampus |  |  |  |  |  |
| IL12 | 1.35 ± 0.35 | 12.36 ± 3.84* | 6.74 ± 1.59 | 2.11 ± 0.43<sup>&</sup> | 10.90 ± 2.29* |
| IL23 | 1.00 ± 0.54 | 10.21 ± 1.39* | 4.48 ± 0.57 | 1.89 ± 0.55<sup>&</sup> | 2.92 ± 0.95 |
| Arg-1 | 6.41 ± 1.07 | 1.41 ± 0.35* | 0.83 ± 0.26* | 6.85 ± 2.89<sup>&</sup> | 1.19 ± 0.41* |

### Cholesterol- and/or BP-lowering therapy affect microglia phenotype in aged ApoE^-/-^ mice

The age-related increase of CD68 (monocyte/macrophage activation marker) in ApoE^-/-^ mice was significantly lower in groups treated with simvastatin (**Fig.3d**). Since a majority of CD68+ cells were positive for ionized calcium-binding adapter molecule-1 (Iba-1; common microglia/macrophage marker) we assessed how age and treatment affected the number of total and CD68+ brain macrophages, and investigated their morphology by counting the numbers of ramified (= resting), intermediate and round (= activated) Iba-1+ cells. Ageing associated with increased numbers of CD68+ Iba-1+ cells in two hippocampus regions that are linked to spontaneous retrieval of episodic contexts and context-dependent retrieval of information, formation of new episodic memories and spontaneous exploration of novel environments (i.e., Cornu Ammonis (CA) 1 and Dentate Gyrus (DG), respectively) but not in the cortex of ApoE^-/-^ mice (**Fig.3e/f & Supplementary Fig.2a**), indicative of region-specific augmentation of macrophage activation. Both hydralazine- and simvastatin-treated mice presented with lower numbers of CD68+ Iba-1+ cells in the CA1 region of the hippocampus (**Fig.3e**); yet, only simvastatin revealed similar responses in the DG (**Fig.3f**). Neither drug affected CD68+ Iba-1+ cell counts in the cortex (**Supplementary Fig.2a**). The total number of Iba-1+ cells significantly increased with age in the CA1 region of the hippocampus (**Fig.3g**) but not in DG or cortex (**Supplementary Fig.2b/c**), suggesting region-specific proliferation of resident macrophages or accumulation of monocyte-derived macrophages. Yet, significant age-related changes in Iba-1+ cell morphology were only observed in the hippocampal DG region as evidenced by a reduced proportion of ramified and increased amount of intermediate microglia (**Table 5**), adding to the notion that age elicits brain region-specific responses in chronically hypercholesterolemic mice. Treatment differently affected overall Iba-1+ cell number and morphology. While hydralazine treatment showed no apparent effects in any of the investigated brain regions (**Table 5**), mice treated with simvastatin presented with a markedly lower overall count of Iba-1+ cells (**Fig.3g**) and significantly reduced percentage of ramified and increased quantity of intermediate Iba-1+ cells compared to aged ApoE^-/-^ mice in the hippocampal CA1 region and in the cortex (**Table 5**). Interestingly, simvastatin-treated mice presented with overall lower numbers of Iba-1+ cells compared to young ApoE^-/-^ in both hippocampal DG and cortex (**Supplementary Fig.2**) and marked changes in Iba-1+ cell morphology in all brain regions (**Table 5**). Similarly, combination treatment significantly reduced the number of ramified cells compared to young ApoE^-/-^ mice in all brain regions investigated but markedly increased the proportion of activated cells compared to aged ApoE^-/-^ in the hippocampal brain region without affecting overall Iba-1+ cell counts (**Table 5**). Macrophages are highly plastic and adopt a variety of phenotypes on the scale between classically activated and alternatively activated phenotypes in response to various stimuli ^29^. Simvastatin treatment attenuated the expression of several markers, characteristic for classical activation (i.e., tumor necrosis factor alpha (TNF-α), IL6, inducible nitric oxide synthase (iNOS); **Table 3**) and significantly mitigated CD86 expression (**Fig.3h**) that has been shown to involve in pro-inflammatory phenotype-mediated retinal degeneration and Alzheimer’s disease (AD) ^30,31^. Of markers typical for alternatively activated macrophages, only Arg-1 was detected in cortical and hippocampal brain tissue in our model. Arg-1 mRNA expression significantly reduced with age in ApoE^-/-^ mice, but aged mice treated with simvastatin presented with higher hippocampal Arg-1 mRNA expression (**Table 4**). Together, these data are suggestive of brain region-specific macrophage activation during ageing in ApoE^-/-^ mice and an alternatively activated phenotype predominance after simvastatin treatment. Lastly, the age-related accumulation of CD3 was diminished in all treatment groups (**Fig.3i**). Using Western blotting, CD3 was not detected in brain tissue of young ApoE mice. This experimental group is therefore not shown in Fig.3I.

**Table 5:**
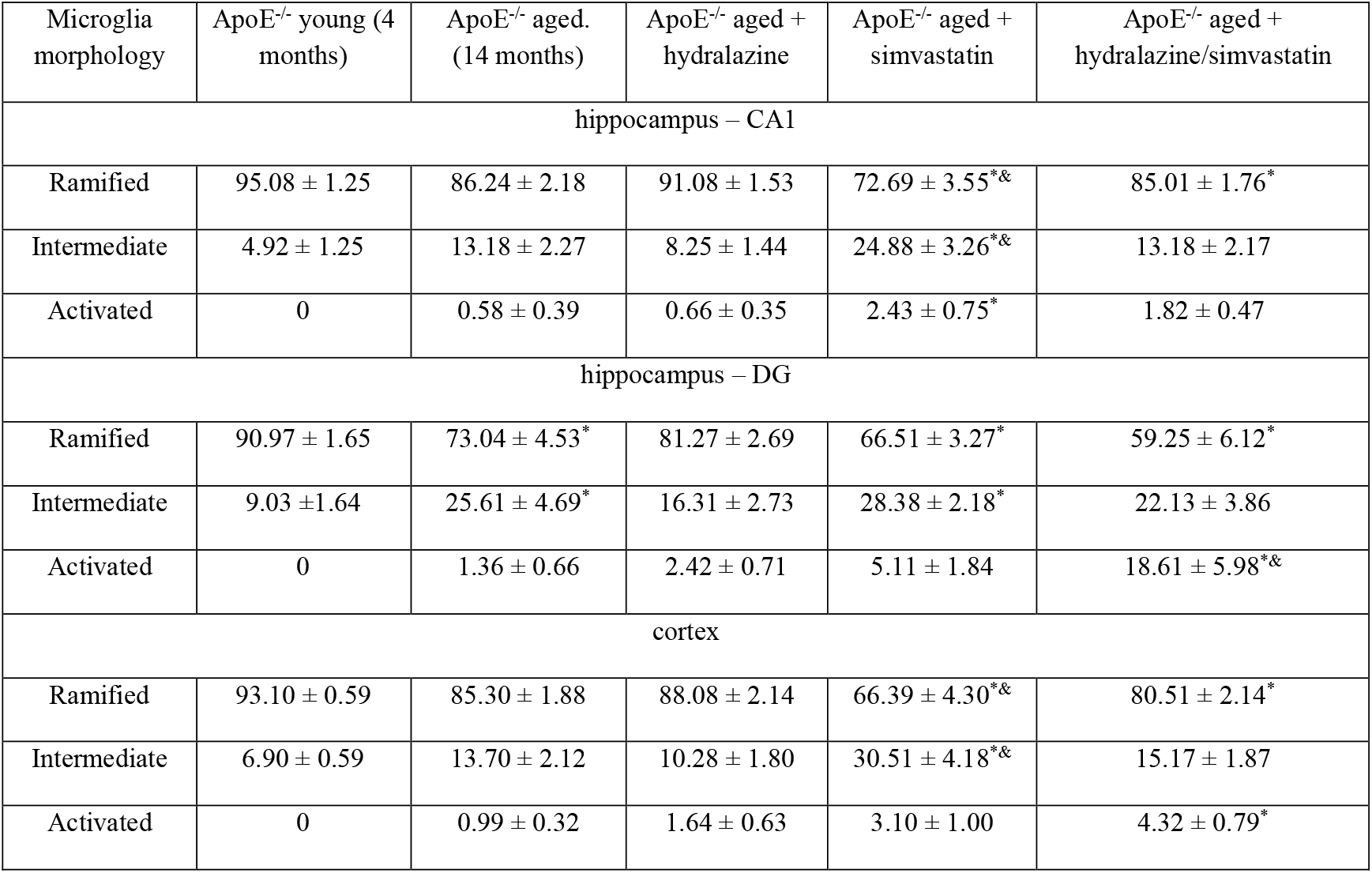
Treatment differently affects microglia morphology in vivo. Morphological quantification of immune-stained Iba-1+ cells in hippocampus regions (CA1 and DG) and cortex classified into *Ramified* (resting state), *Intermediate* and *Round* (active state). ApoE – apolipoprotein E, CA – cornu ammonis, DG – dentate gyrus, Iba-1 – ionized calcium-binding adapter molecule-1. *Values are expressed as mean (of all Iba-1+ cells per type) +/- SEM of N=3 tissues samples for young and aged + combination treatment; N=4 tissues samples for aged control and simvastatin treatment; N=5 tissues samples for hydralazine treatment where N denotes number of independent biological replicates. For CA1 and DG, * denotes P ≤ 0*.*05 relative to young control, & denotes P ≤ 0*.*05 relative to aged control after one-way ANOVA followed by Tukey’s post hoc testing; for cortex, * denotes P ≤ 0*.*05 relative to young control, & denotes P ≤ 0*.*05 relative to aged control after Kruskal Wallis followed by Dunn’s post hoc testing*.

| Microglia morphology | ApoE<sup>-/-</sup> young (4 months) | ApoE<sup>-/-</sup> aged. (14 months) | ApoE<sup>-/-</sup> aged + hydralazine | ApoE<sup>-/-</sup> aged + simvastatin | ApoE<sup>-/-</sup> aged + hydralazine/simvastatin |
| --- | --- | --- | --- | --- | --- |
| hippocampus – CA1 |  |  |  |  |  |
| Ramified | 95.08 ± 1.25 | 86.24 ± 2.18 | 91.08 ± 1.53 | 72.69 ± 3.55*& | 85.01 ± 1.76* |
| Intermediate | 4.92 ± 1.25 | 13.18 ± 2.27 | 8.25 ± 1.44 | 24.88 ± 3.26*& | 13.18 ± 2.17 |
| Activated | 0 | 0.58 ± 0.39 | 0.66 ± 0.35 | 2.43 ± 0.75* | 1.82 ± 0.47 |
| hippocampus – DG |  |  |  |  |  |
| Ramified | 90.97 ± 1.65 | 73.04 ± 4.53* | 81.27 ± 2.69 | 66.51 ± 3.27* | 59.25 ± 6.12* |
| Intermediate | 9.03 ± 1.64 | 25.61 ± 4.69* | 16.31 ± 2.73 | 28.38 ± 2.18* | 22.13 ± 3.86 |
| Activated | 0 | 1.36 ± 0.66 | 2.42 ± 0.71 | 5.11 ± 1.84 | 18.61 ± 5.98*& |
| cortex |  |  |  |  |  |
| Ramified | 93.10 ± 0.59 | 85.30 ± 1.88 | 88.08 ± 2.14 | 66.39 ± 4.30*& | 80.51 ± 2.14* |
| Intermediate | 6.90 ± 0.59 | 13.70 ± 2.12 | 10.28 ± 1.80 | 30.51 ± 4.18*& | 15.17 ± 1.87 |
| Activated | 0 | 0.99 ± 0.32 | 1.64 ± 0.63 | 3.10 ± 1.00 | 4.32 ± 0.79* |

### Cholesterol- and/or BP-lowering therapy attenuate memory deficits in aged ApoE^-/-^ mice

In an open field test, only simvastatin treatment significantly improved age-related impaired mobility in ApoE^-/-^ mice (**Fig.4a**). Combination treatment with hydralazine revealed no apparent effects on mobility; yet, statistical analysis disclosed no difference to young control. Likewise, hippocampal-dependent memory function (**Fig.4b**) and spatial short-term memory function (**Fig.4c**) significantly improved in all treatment groups containing simvastatin. Brain-derived neurotrophic factor (BDNF), an important regulator of white matter integrity, hippocampal long-term potentiation and consequently, learning and memory ^32^, was negatively affected by age in ApoE^-/-^ mice, and only simvastatin treatment mitigated the age-related reduction of hippocampal BDNF expression (**Fig.4d**). Similarly, immune staining of hippocampal BDNF and the neuronal marker NeuN revealed evident differences between young and aged mice (**Supplementary Table 3**, representative images shown in **Fig.4e**). When normalized to the number of NeuN+ cells, BDNF expression was significantly lower in aged brains compared to young ApoE^-/-^ mice, and only significantly higher after treatment with simvastatin alone (**Fig.4e**). NeuN immunostaining, however, was significantly higher in all treatment groups that received simvastatin compared to untreated aged ApoE^-/-^ mice (**Supplementary Table 3**). Similar to the hippocampus, immune staining for BDNF in the cortex revealed significant differences between young and aged ApoE^-/-^ mice, but all treatment regimens presented with high BDNF protein expression in this brain region (**Supplementary Table 3**). Most notably, however, NeuN immune staining only significantly differed in hippocampus but not cortex where neither age nor treatment affected NeuN expression (**Supplementary Table 3**), further supporting a prominent hippocampus-dependent memory impairment in aged chronically hypercholesterolemic ApoE^-/-^ mice.

**Figure 4:**
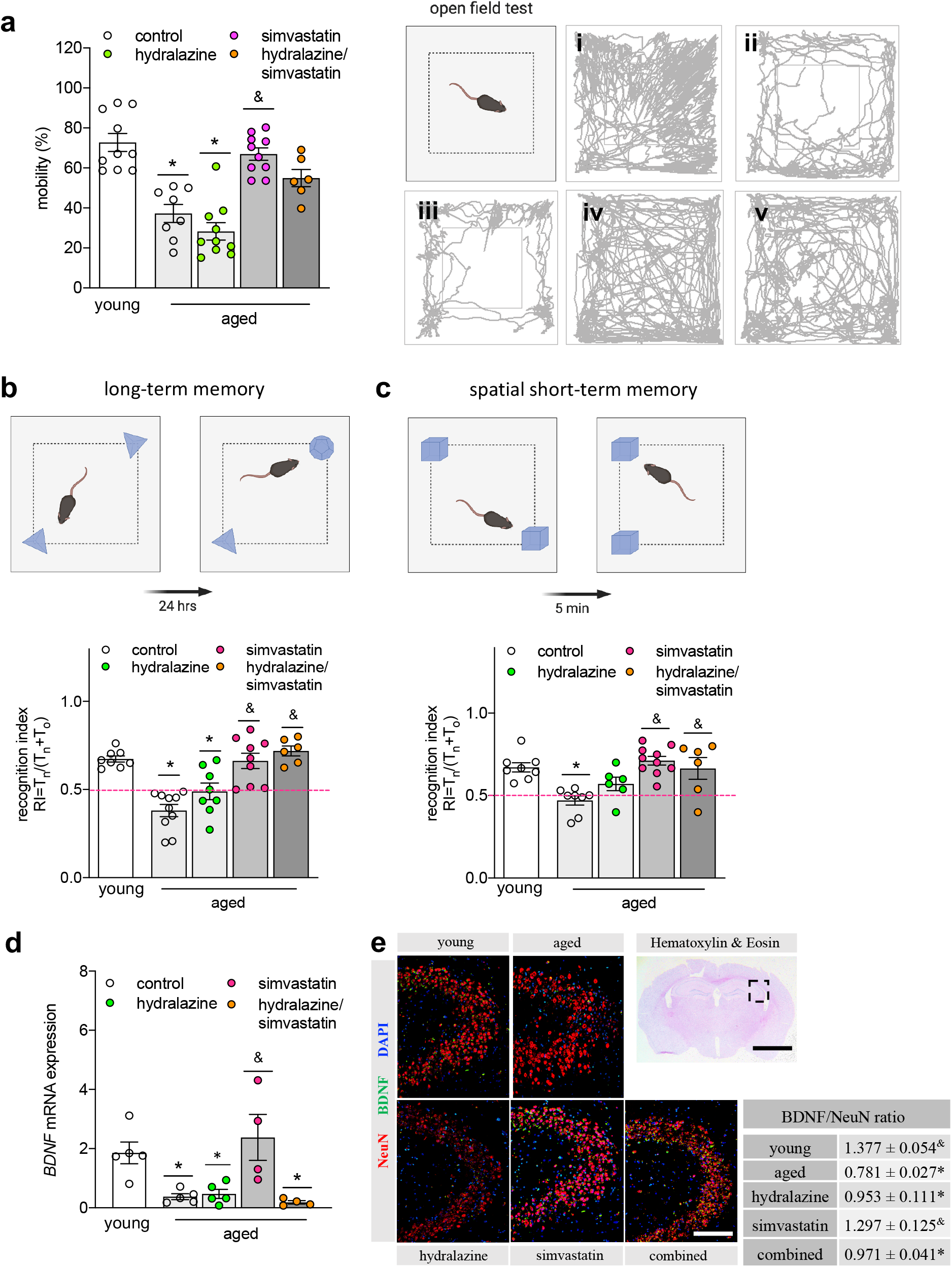
Simvastatin mitigates impaired memory function of aged ApoE^-/-^ mice. **(a)** Quantification of open field test assessing mobility in aged ApoE^-/-^ mice treated with hydralazine (green circles), simvastatin (pink circles) and a combination treatment (orange circles) in comparison to young and aged control mice (open circles). (N=10 per group; some animals were excluded due to total mobile times below 20s). Representative images showing young control, aged control, aged + hydralazine, aged + simvastatin, and aged + hydralazine/simvastatin combined. **(b)** Effect of lipid- and BP-lowering treatment on long-term memory function, and on **(c)** spatial long-term memory function. (N=10 per group; some animals were excluded due to total object exploration times below 20s). The pink dotted line indicates the point where the animal has no preference for either object. **(d)** Effect of lipid- and BP-lowering treatment on hippocampal BDNF mRNA expression (N=5 for young, aged and hydralazine-treated groups; N=4 for simvastatin- and combination-treated groups). **(e)** Representative images of BDNF (green) and NeuN (red) immune staining (scale bar 50 μm) of the hippocampal CA region (region indicated in Hematoxylin-Eosin-stained coronal brain section; scale bar 25mm) in young, aged control, aged + hydralazine, aged + simvastatin, and aged + combined treatment. Quantification of BDNF-NeuN ratio comparing young and aged ApoE^-/-^ mice with treatment groups (N=3 for young and aged + combination treatment; N=4 for aged control and simvastatin treatment; N=5 for hydralazine treatment). ApoE – apolipoprotein E, BDNF – brain-derived neurotrophic factor, BP – blood pressure, CA – cornu ammonis, RI – recognition index. Data expressed as mean ± SEM. N denotes number of independent biological replicates. In (**a)**, * denotes P ≤ 0.05 relative to young control, & denotes P ≤ 0.05 relative to aged control after Kruskal-Wallis followed by Dunn’s *post hoc* testing. In (**b)-(d)**, * denotes P ≤ 0.05 relative to young control, & denotes P ≤ 0.05 relative to aged control after one-way ANOVA followed by Tukey’s *post hoc* testing. All schematics were created with BioRender.

### Simvastatin and hydralazine exert cell-specific effects

The observed differences of simvastatin and hydralazine treatment on inflammatory responses *in vivo*, led us to evaluate their effects *in vitro* using different immune cell subtypes of human origin. Utilizing a human monocytic cell-line (THP-1) allowed us to assess drug effects not only in monocytes but also in monocyte-derived macrophages after differentiation. Flow cytometric analyses of monocytic or phorbol 12-myristate 13-acetate (PMA)-differentiated THP-1 cells revealed a strong pro-inflammatory effect of hydralazine in monocytic cells (i.e., augmentation of Lipopolysaccharide (LPS)-associated CD14+ CD16+ surface expression; 2.5-fold; **Fig.5a**) compared to PMA-differentiated THP-1 cells where hydralazine did not increase CD14+ CD16+ surface expression **(Fig.5b**). Similar cell type-specific effects were observed for the surface expression of the activation marker CD69 that significantly increased with hydralazine but not simvastatin treatment in monocytic THP-1 cells but not in PMA-differentiated macrophages (**Supplementary Table 4**). Pro-inflammatory cytokine profiling in THP-1 cells revealed that LPS-induced augmentation of intracellular IL6 protein abundance was not affected by simvastatin treatment but exacerbated in the presence of hydralazine (**Fig.5c**). Hydralazine alone (without LPS pre-stimulation) disclosed similar pro-inflammatory effects (**Fig.5d**). Different from monocytic cells, LPS-induced IL6 expression was similarly reduced in the presence of simvastatin or hydralazine in PMA-differentiated THP-1 cells (**Fig.5e**), indicative of cell type-specific drug responses. Similarly, LPS-induced augmentation of TNF-α mRNA expression only reversed in simvastatin-containing treatment groups but not with hydralazine in monocytic THP-1 cells (**Fig.5c**). In support of this, LPS-induced elevation of TNF-α and IL6 secretion in murine bone-marrow-derived macrophages (BMDMs) reduced with both simvastatin and hydralazine treatment (**Fig.5g/h**). Moreover, we detected lower surface expression of activation markers (i.e., CD86, CD80) in BMDMs of all treatment groups (**Supplementary Fig.3a/b**). Lastly, we used primary human monocytes and monocyte-derived macrophages to confirm cell type-specific drug effects: hydralazine significantly induced cell activation (i.e., increased CD69 expression) that affected monocytes with a higher magnitude than macrophages (**Fig.5i, Supplementary Table 4**) while simvastatin presented with no appreciable effects on CD69 expression (**Fig.5k, Supplementary Table 4**). Similar to CD69, hydralazine increased monocyte and macrophage CD3 expression in both primary human cells and THP-1 cells (**Supplementary Fig.4a-e, Supplementary Table 4**), which has been shown to involve in the delivery of pro-inflammatory cytokines (i.e., TNF-α) by macrophages.^33^ Notably, none of the drugs showed effects on T-cell activation (**Supplementary Fig.5**). Together, these data suggest cell type-specific drug effects that may benefit simvastatin as it exerted anti-inflammatory effects in both activated monocytic cells and activated macrophages. Hydralazine on the other hand, induced pro-inflammatory signatures especially in monocytic cells with potential implications for conditions characterized by elevated monocyte counts like chronic hypercholesterolemia, which may explain some of the effects we observe *in vivo*.

**Figure 5:**
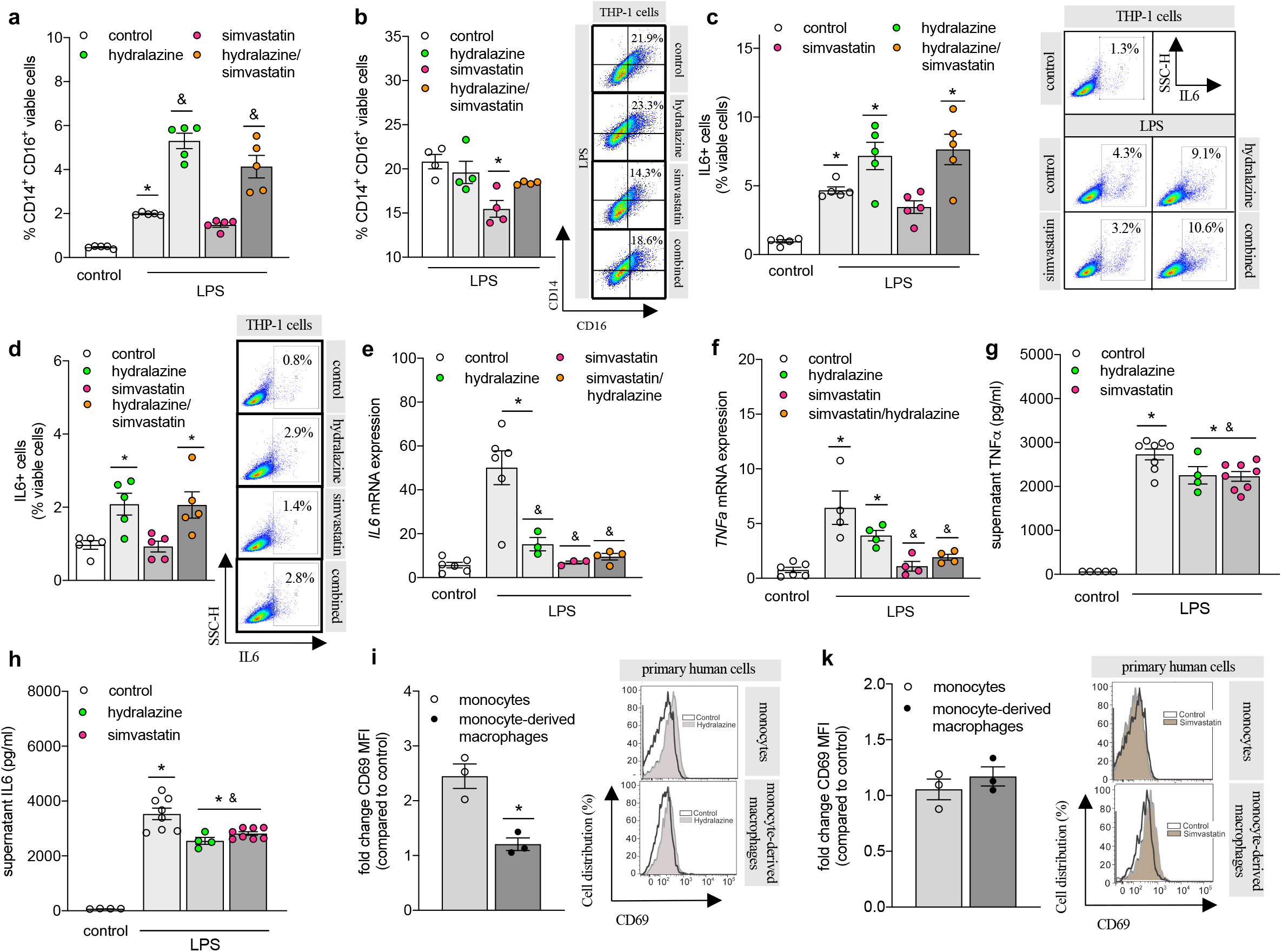
Simvastatin reduces monocyte activation. **(a)** Flow cytometry analysis of CD14+ CD16+ cells in monocytic THP-1 cells (N=5 per group) and (**b**) PMA-differentiated THP-1 cells (N=4 per group) following LPS activation and treatment with simvastatin, hydralazine or a combination of both (hydralazine – green circle, simvastatin – pink circles, combination – orange circles). Representative dot plots showing typical CD14-CD16 surface expression. **(c)** Effect of hydralazine and simvastatin on LPS-induced intracellular IL6 expression in human monocytic THP-1 cells determined by flow cytometry (N=5 per group). Representative dot plot showing the typical IL6+ cell distribution. **(d)** Effect of hydralazine and simvastatin on intracellular IL6 expression in human monocytic THP-1 cells determined by flow cytometry (N=5 per group). Representative dot plot showing the typical IL6+ cell distribution. **(e)** Effect of hydralazine and simvastatin on IL6 mRNA expression in human THP-1 cells PMA-differentiated to macrophages determined by qPCR (N=3-6 per group). (**f**) Effect of hydralazine and simvastatin on LPS-induced TNF-α mRNA expression in human monocytic THP-1 cells determined by qPCR (N=6 in control and combination-treated groups; N=4 in LPS control, hydralazine- and simvastatin-treated groups). **(g)** Effect of hydralazine and simvastatin on LPS-mediated TNF-α and **(h)** IL6 release in murine bone marrow-derived macrophages assessed by ELISA (N=4 for untreated control and hydralazine-treated cells, N=8 for LPS-treated control and simvastatin-treated cells). (**i**) Degree of myeloid cell activation in response to hydralazine determined by flow cytometric assessment of CD69 surface expression (expressed in fold change compared to control; non-normalized data can be found in **Supplementary Table 4**) in human primary monocytes and monocyte-derived macrophages (N=3 per group). Representative histograms showing the typical CD69+ cell distribution. **(k)** Degree of myeloid cell activation in response to simvastatin determined by flow cytometric assessment of CD69 surface expression (expressed in fold change compared to control; non-normalized data can be found in **Supplementary Table 4**) in human primary monocytes and monocyte-derived macrophages (N=3 per group). Representative histograms showing the typical CD69+ cell distribution. IL – interleukin, LPS – lipopolysaccharide, PMA – phorbol 12-myristate 13-acetate, TNF-α – tumor necrosis factor alpha. Data expressed as mean ± SEM. N denotes number of independent biological replicates. In **(a)-(h)** * denotes P ≤ 0.05 relative to control, & denotes P ≤ 0.05 relative to LPS-treated control after one-way ANOVA followed by Tukey’s *post hoc* testing; in **(i)**, * denotes P ≤ 0.05 relative to control for unpaired t-test.

## Discussion

The present study provides compelling evidence that chronic hypercholesterolemia majorly contributes to the development of memory impairment, involving pro-inflammatory processes. The herein established link between chronically elevated plasma cholesterol and the development of memory impairment in aged mice is based on findings, showing that only early exposure to elevated plasma cholesterol (i.e., as evident in ApoE^-/-^ mice already at 4 months of age), indicative of chronic hypercholesterolemia, results in reduced long-term and spatial memory function at an older age (i.e., 12 months of age). This conclusion aligns with data obtained in epidemiological studies supporting a correlation between early- or mid-life cholesterol levels to dementia later in life ^34,35^. Similar to our findings obtained in aged normocholesterolemic WT mice, no association between cognitive function and cholesterol levels was found in studies testing older patients ^36,37^. Interestingly, it has been shown that female ApoE^-/-^ mice display greater cognitive impairment than male ApoE^-/-^ mice ^38,39^. The fact that female ApoE^-/-^mice develop more severe atherosclerosis and vascular impairment earlier ^40^ supports the hypothesis that atherosclerosis contributes to cognitive dysfunction in this model. This is further signified by findings showing an absence of cognitive deficits in female brain-specific ApoE^-/-^ mice that are characterized by absence of systemic hypercholesterolemia ^39^. Thus, further investigations of sex-specific hypercholesterolemia effects on systemic and neuro-inflammation are needed.

Although hypercholesterolemia was generally accompanied by higher-than-normal BP in aged ApoE^-/-^ mice, cholesterol-lowering rather than BP-lowering therapy significantly improved the impaired spatial memory of aged ApoE^-/-^ mice. This further positions early life exposure to high cholesterol levels as major contributor to cognitive dysfunction in addition to potential BP-mediated structural and functional alterations in the brain ^3^. Unlike statins for dyslipidemia treatment, hydralazine is not the first line medication for treating elevated BP. Nonetheless, hydralazine appears as first choice of BP-lowering therapy in numerous experimental studies ^3,41,42^, and is often used in the clinic to treat preeclampsia. Besides its vasorelaxant properties, it is thought to exert effects on the immune system, specifically on T-cell transmigration ^23^ and apoptosis ^43^. However, hydralazine-induced increases of cellular CD3-zeta chain content in T-cells are discussed to facilitate excessive drug-induced auto-immunity ^44^. The pronounced enhancement of monocytic cell activation we observed in the presence of hydralazine might be resultant from reactive metabolites deriving from hydralazine oxidation ^45^ or from hydralazine-mediated inhibition of DNA methyltransferase ^46^, which are postulated as major mechanisms in drug-induced auto-immunity, might limit its potency to resolve neuro-inflammatory processes and thus, improve memory deficits in our model. Moreover, it might limit therapeutic efficacy of other drugs when administered in combination. This effect calls for the need to test for potential interactions between BP-lowering drugs of different nature and statins for potential consequences on immune regulation and cognitive impairment. It will be of utmost importance to further investigate the value of different classes of BP-lowering drugs in multi-morbid systems.

The activation of the immune system emerged as an important driver in the aged and vulnerable brain. Experimental studies verified the significance of inflammation in CVD progression ^47,48^ and showed that inflammation associated with cardiovascular risk factors or CVD negatively affects cognitive function ^17,18,49,50^. However, precise pro-inflammatory mechanisms contributing to accelerated cognitive decline and dementia risk require further elucidation. Similar to a study that links the Th17 - IL17 pathway to high dietary salt-induced cognitive impairment in mice ^50^, we observe elevated IL17 plasma levels in cognitively impaired aged ApoE^-/-^ mice with elevated cholesterol and BP levels. Further, the link between circulating IL17 levels and memory performance is strengthened as simvastatin-mediated lowering of plasma cholesterol, BP and IL17 levels improved memory function. Another experimental study convincingly associates high fat diet-induced hypercholesterolemia in aged ApoE^-/-^ mice not only with elevated plasma levels of IL6, TNF-α, and interferon (IFN)-γ but also higher transcript levels of pro-inflammatory chemokine MCP-1 in brain tissue of these mice ^51^. In our study, we witness a similar increase of such key chemokines responsible for regulating migration and infiltration of monocytes in brain tissue of standard chow-fed aged ApoE^-/-^ but not aged-matched WT mice. Consequently, the accumulation of pro-inflammatory Ly6Chi monocytes in the brain was absent in aged WT mice where plasma cholesterol levels only increase later in life and thus, positions pro-inflammatory monocytes as important contributors to neuro-inflammation emanating from chronic hypercholesterolemia. The concept of hypercholesterolemia-associated neuro-inflammation was recently investigated in aged hypercholesterolemic ApoE^-/-^ mice where choroid plexus lipid accumulation induced leukocyte infiltration that extended to the adjacent brain parenchyma, demonstrating that lipid-triggered complement cascade activation promotes leukocyte infiltration into the choroid plexus with implications for immune system activation, cognitive decline in late onset AD and atherosclerotic plaque formation ^52^. Together with the finding that brain-specific ApoE deficiency protects from cognitive decline in the absence of hypercholesterolemia ^39^, this further signifies the importance of chronic hypercholesterolemia in neuro-inflammatory and -degenerative processes observed in our model. The herein characterized brain region-specific neuro-inflammation (i.e., brain macrophage proliferation and activation, cytokine and classical/alternative microglia activation marker expression) together with distinct brain region-specific changes after cholesterol-lowering therapy, however, also suggest additional simvastatin-mediated mechanisms independent of cholesterol-lowering.

Our experimental strategy demonstrates that simvastatin therapy proves effective in reducing neuro-inflammation associated with chronic hypercholesterolemia in aged ApoE^-/-^ mice. In particular, its beneficial hippocampal-specific effects on Iba-1+ cell number and morphology, CD86 and Arg-1 expression in our model point towards additional therapeutic mechanisms independent of cholesterol-lowering. Simvastatin’s anti-inflammatory character might be resultant from its potency to regulating proliferation and activation of monocytes and macrophages ^19,21^, and to modulating myeloid cell phenotype ^53^ and secretory function ^54,55^. The presented *in vitro* findings describe simvastatin’s direct therapeutic effect on monocyte and macrophage activation and extent previous reports that showed an effective reduction of cytokine secretion in PMA-differentiated THP-1 cells pre-treated with simvastatin ^54^. In its role as modulator for IL secretion from monocytes and macrophages, simvastatin was shown to indirectly inhibit IL17 secretion from CD4+ T-cells ^56^. Moreover, simvastatin has been shown to directly inhibit the expression of a transcription factor responsible for controlling IL17 production in CD4+ T-cells ^56^. These findings support our herein presented data showing a reduction of IL17 plasma levels *in vivo* only after simvastatin treatment. However, it remains to be determined whether the simvastatin effects in our model are mediated through direct modulation of Th17 responses, indirectly through modulation of myeloid cell activation or primarily through its lipid-lowering capacity. Because simvastatin has also been associated with Th2 immune response and neuronal recovery ^57,58^ we cannot exclude a potential therapeutic effect on T-cells in our model. Nonetheless, our results point to therapeutic effects through monocyte modulation, which aligns with studies conducted in patients at high risk for vascular events where a simvastatin-induced down-regulation of the angiotensin II type 1 receptor on monocytes but not T-cells significantly affected angiotensin II activity and thus, contributes to its anti-inflammatory profile and therapeutic effects in CVD ^59^. Moreover, in patients with hypercholesterolemia, simvastatin was shown to reduce monocyte secretory function ^55,60^. Most interestingly in respect to neurodegeneration is a study reporting a lowering of the CD14+CD16+ “intermediate” monocyte subset in purified monocytes isolated from peripheral blood of human immunodeficiency virus (HIV) patients in response to an *ex vivo* treatment with simvastatin, as this particular subset of monocytes is closely linked to HIV-associated neurocognitive disorders^53^.

In contrast to case reports discussing adverse effects of statins on cognitive function ^12,13^, our study describes favorable outcomes for memory function in aged ApoE^-/-^ mice after statin therapy, supporting findings from two large randomized control trials on simvastatin ^61^ and pravastatin ^62^ where no link between statin use and cognitive decline was observed. Although lipophilic statins, such as simvastatin are thought to cross the blood– brain barrier, leading to a reduction of cholesterol availability and thereby, disturbing the integrity of the neuronal and glial cell membrane ^63^, simvastatin treatment had no negative effects on hippocampal neurons, and significantly increased hippocampal and cortical BDNF expression in our model. Most beneficial effects of simvastatin in the brain are thought to root from the promotion of hippocampal neurogenesis, the inhibition of mesangial cell apoptosis ^64,65,^ the enhancement of neurotrophic factors, and the restriction of inflammation ^66^. In light of this, simvastatin treatment markedly increased hippocampal expression of BDNF, which has been linked to improved functional recovery after stroke and the amelioration of depressive-like behavior ^67,68^, and may underlie the improvements in hippocampal-dependent memory function we observe in our model. Besides, preventative simvastatin treatment resulted in lower Aβ 40/42 levels in cerebrospinal fluid of AD mice, attenuated neuronal apoptosis and improved cognitive competence of the hippocampal network ^69^, suggesting that statin treatment may prevent the age-related, progressive neuropathy characteristic for 3xTg-AD mice. Further investigation of Amyloid β accumulation in respect to neuro-inflammation and its modifiability by cholesterol-lowering or anti-inflammatory therapies would certainly elevate the impact of statin therapy beyond the field of CVD and associated target organ damage.

Taken together, our study convincingly links chronic hypercholesterolemia to myeloid cell activation, neuro-inflammation and memory impairment, supporting the notion that early rather than late-life exposure to cardiovascular risk factors promotes the development of cognitive dysfunction. Cholesterol-lowering therapy provides effectiveness to improving memory function potentially by reducing monocyte-driven inflammatory events and hence, emerges as safe therapeutic strategy to control CVD-induced memory impairment and represents another benefit of statins in patients with hypercholesterolemia.

### Study limitations and outlook

Controversies regarding BP phenotype in ApoE^-/-^ mice persist, however, it is difficult to draw conclusions from the current literature as studies differ in duration, diet and animal age. Dinh and colleagues reported a lack of BP elevation in aged ApoE^-/-^ mice subjected to high-fat diet compared to WT mice on conventional diet.^51^ Another study reports that high-fat diet increases BP already after 12 weeks.^70^ In contrast to the herein presented results, an earlier study suggested no differences in BP between aged ApoE^-/-^ and WT mice on normal chow diet (140 ± 7.6 mmHg in ApoE^-/-^ mice and 136 ± 7.4 mmHg in WT mice).^71^ The study, however, did not assess BP differences between young and aged ApoE^-/-^ mice. Considering the current knowledge in the field, it remains to be further investigated whether or not changes in systemic BP contribute to the inflammatory and the brain phenotype described in this model. Furthermore, neither memory impairment nor neuro-inflammation in our study were assessed longitudinally. However, our data convincingly show apparent memory deficits and obvious signs of neuro-inflammation and -degeneration in ApoE^-/-^ mice at the age of 12 months, which goes in line with findings of previously published studies.^38,39^ It is therefore likely that the applied treatment regimens not only attenuate but also reverse brain degeneration in aged ApoE^-/-^ mice. In order to decipher cell type-specific treatment effects on circulating immune cells, systemic assessment of different immune-related tissue compartments would be necessary. Moreover, understanding the mechanism of action by which simvastatin or other CVD therapeutics influence neuro-inflammatory events and affect monocyte and macrophage polarization *in vivo* will require further investigation using more advanced single-cell techniques, which have recently provided important insight into the plasticity of these cell types and the existence of a wide stimuli-driven network of myeloid cell behavior^72,73^ that may drastically impact neurological performance. In line with that, it would be of interest how cholesterol- and BP-lowering therapies affect atherosclerosis progression (i.e., lesion size and plaque stability) in this model and to what extent this affects memory functions. Although statins are thought to cause regression of atherosclerosis in humans, it has been shown that the amount of lesion regression is small compared with the magnitude of the observed clinical benefit. Similarly, preclinical studies also showed beneficial statin effects on plaque stability in ApoE^-/-^ mice,^74,75^ however, there seems to be an ongoing debate about their ability to actually reduce lesion size. Lastly, sex-specific differences need to be investigated as our study was solely performed in male mice. Especially since previous studies convincingly showed an increased susceptibility of female mice to the development of cognitive impairment in response to chronic hypercholesterolemia,^38,39^ treatment responses may also differ sex-specifically.

## Methods

### Materials

All chemical reagents and solutions were purchased from Fisher Scientific (Göteborg, Sweden), Saveen & Werner (Limhamn, Sweden) or Sigma-Aldrich (Stockholm, Sweden) unless otherwise stated. Commercially available primary antibodies against CD3 (Bio-techne, UK), PSD-95, SNAP-25, NeuN and BDNF (Abcam, UK), CD68 (Invitrogen, Sweden) and Iba-1 (Wako, Japan) were used for immunofluorescence. Secondary antibodies Alexa Fluor 488 donkey anti-mouse, anti-rabbit, goat anti-rabbit or goat anti-rat Fluor 594 (Nordic Biosite, Sweden) were used for visualization. Primers for qPCR were purchased from Eurofins (Ebersberg, Germany).

### Animals

This investigation conforms to the Guide for Care and Use of Laboratory Animals published by the European Union (**Directive 2010/63/EU**) and with the ARRIVE guidelines. All animal care and experimental protocols were approved by the institutional animal ethics committees at the University of Barcelona (CEEA) and Lund University (5.8.18-12657/2017) and conducted in accordance with European animal protection laws. Male wild-type (WT) C57Bl/6J mice and Apolipoprotein E knockout (ApoE^-/-^) mice (B6.129P2-Apoe (tm1Unc)/J) were obtained from Jackson Laboratories and bred in a conventional animal facility under standard conditions with a 12h:12h light-dark cycle, and access to food (standard rodent diet) and water *ad libitum*. Mice with a body weight BW ≥ 25g were housed in groups of 4-5 in conventional transparent polycarbonate cages with filter tops. ApoE^-/-^ and C57Bl/6J (WT) mice at the age of 4 and 12 months (N=10 per group) were used to assess the effect of ageing on inflammatory status and brain phenotype (**Fig.1a**). At the age of 12 months, another group of ApoE^-/-^ mice was randomly assigned to the following experimental groups using the computer software *Research Randomizer* (http://www.randomizer.org/): aged control, aged + hydralazine (25 mg/L), aged + simvastatin (2.5 mg/L), and aged + hydralazine/simvastatin combination (N=10 for simvastatin and hydralazine treatment; N=8 for combination treatment). The calculated doses given to the mice were 5 mg/kg/d for hydralazine and 0.5 mg/kg/d for simvastatin, which corresponds to 10 mg/kg/d and 20 mg/kg/d in humans, respectively (allometric scaling was used to convert doses amongst species ^76^). Treatment was administered via drinking water over a course of two months. The mice in this experimental cohort were 14 months of age at time of sacrifice (**Fig.2b**). ApoE^-/-^ mice at the age of 4 months were used as young controls (N=10). To ensure blinding, experiments were performed after the animals and samples had received codes that did not reveal the identity of the treatment. In order to obey the rules for animal welfare, we designed experimental groups in a way that minimizes stress for the animals and guarantees maximal information using the lowest group size possible when calculated with a type I error rate of α = 0.05 (5%) and Power of 1-β > 0.8 (80%) based on preliminary experiments. At termination, brain tissue was distributed to the different experiments as follows: Young (4 months) and aged (12 months) WT and ApoE^-/-^ mice (N=10 each) for flow cytometric assessment of one hemisphere, N=5 each for cortex and hippocampus fractionation of one hemisphere and N=5 for whole hemisphere RNA and protein isolation (**Fig.1a**). Young (4 months), aged (14 months), hydralazine- and simvastatin treated (N=5 each) and combination-treated (N=3) ApoE^-/-^ mice for whole brain cryo-sectioning, cortex and hippocampus fractionation of one hemisphere (N=5 each) as well as one hemisphere (N=5 each) for whole hemisphere RNA and protein isolation (**Fig.2b**).

### Open Field testing

To test novel environment exploration, general locomotor activity, and screen for anxiety-related behavior mice were subjected to an open field exploration task using a video tracking system and the computer software EthoVision XT® (Noldus Information Technology, Netherlands) as described before^.17^ Mice were placed individually into an arena (56 × 56 cm), which was divided into a grid of equally sized areas. The software recorded each line crossing as one unit of exploratory activity. The following behavioral parameters were measured: activity, active time, mobile time, slow activity, mobile counts, and slow mobile counts.

### Novel object recognition (NOR)

As previously described,^3,17,77^ a NOR task was employed to assess non-spatial memory components. Briefly, mice were habituated to the testing arena for 10 minutes over a period of 3 days. On test day, each mouse was exposed to two objects for 10 minutes. 5 min or 24 hrs later, mice were re-exposed to one object from the original test pair and to a novel object. The movements of the animal were video tracked with the computer software EthoVision XT® (Noldus Information Technology, Netherlands). A delay interval of 5 minutes was chosen to test short-term retention of object familiarity, and with a delay interval longer of 24 hrs, we tested long-term hippocampus-dependent memory function.^3,78^

### Object placement task

To test spatial recognition memory, mice were placed individually into an arena (56 × 56 cm). Each mouse was exposed to two objects for 10 minutes. 5 min later, mice were re-exposed to both objects, of which one remained in the original place and the second object was moved to a novel place. Exploration of the objects was assessed manually with a stopwatch when mice sniffed, whisked, or looked at the objects from no more than 1 cm away. The time spent exploring the objects in new (novel) and old (familiar) locations was recorded during 5 min.

### Blood pressure measurements

BP was measured in conscious mice using tail-cuff plethysmography (Kent Scientific, CODA, UK). After a one-week handling period, mice were acclimatized to the restrainers and the tail cuff for a training period of 7 days. Data were recorded once mice presented with stable readings over the course of one week. During BP measurements, 30 inflation cycles were recorded of which the first 14 were regarded as acclimatization cycles and only cycles 15-30 were included in the analyses.

### Fluorescence activated cell sorting

Whole blood from mice was collected in EDTA-coated tubes and red blood cells were lysed before samples were incubated in F_c_ block solution followed by primary antibodies. After washing and centrifugation, the supernatant was decanted, and pellets were re-suspended in FACS buffer. Brain tissue was enzymatically digested and homogenized. After density separation using Percoll (GE Healthcare), pellets were reconstituted in F_c_ block prior to staining with antibodies (**Supplementary Table 5**). Data acquisition was carried out in a BD LSR Fortessa cytometer using FacsDiva software Vision 8.0 (BD Biosciences). Data analysis was performed with FlowJo software (version 10, TreeStar Inc., USA). Cells were plotted on forward versus side scatter and single cells were gated on FSC-A versus FSC-H linearity.

### Histology and Immunofluorescence

Coronal brain sections (10 μm thickness) were incubated with antibody against NeuN, BDNF, Iba-1 and CD68 (**Supplementary Table 6**). After several washes, sections were incubated with secondary Alexa Fluor-coupled antibodies (Life Technologies, Sweden) at room temperature. Sections were then mounted in aqueous DAPI-containing media (DapiMount, Life Technologies, Sweden). The specificity of the immune staining was verified by omission of the primary antibody and processed as above, which abolished the fluorescence signal.

### Quantification of immune-histological experiments

For quantification of hippocampal NeuN+ cells, three regions of interest (ROI) in the cortex and three brain sections representing the hippocampus were selected. The number of NeuN+ cells in the pyramidal layer of the CA3 region, of the CA1 region and the granule cell layer of the DG as well as in the cortical ROIs was determined using ImageJ software (ImageJ 1.48v, http://imagej.nih.gov/ij). First, a binary image was generated from the original images. To measure the total number of stained cells the analyze particle feature was chosen and the threshold was maintained at the level that is automatically provided by the program, no size filter was applied. BDNF immune staining in the same three brain regions was quantified, and by dividing the obtained value by the corresponding number of NeuN+ cells the ratio of BDNF per NeuN+ cell was obtained for the respective area. Staining was analyzed in three images per area and mouse. For each experimental group *N=5*. Values are expressed as mean +/- SEM.

### Brain macrophage analysis

Coronal brain sections (10μm thickness) of each experimental group were immune-stained with Iba-1 (WAKO, Japan) and CD68 (Invitrogen, Sweden). The total number of Iba-1+ cells was counted in two regions of the hippocampus (dentate gyrus; DG and the cornu ammonis 1; CA1) in each hemisphere and in five ROIs of the cortex using a fluorescence microscope (Axio Imager .M2, Zeiss). The cells were classified into three activation state groups based on morphology where *Ramified* (resting state) denotes Iba-1+ cells with a small cell body and extensive branched processes, *Intermediate* denotes Iba-1+ cells with enlarged cell body and thickened, reduced branches not longer than twice the cell body length and *Round* (active state) denotes Iba-1+ cells with round cell body without visible branches. In all sections, the total numbers of CD68+ Iba-1+ cells were recorded.

### Western Blotting

For protein isolation, tissue lysates were prepared using RIPA buffer containing 10mM Tris (pH 8.0), 1mM EDTA, 1% Triton-X, 0.1% sodium-deoxycholate, 0.1% SDS, 140mM NaCl and 25μg/ml protease inhibitor cocktail. Lysates underwent five freeze-thaw cycles using liquid nitrogen. The resulting lysates were centrifuged for 30 min at 15,000g and 4°C to remove insoluble material. Protein concentration was determined by PIERCE BCA protein assay. Western blotting was carried out according to standard protocols. Briefly, samples were heated for 10 min at 95°C in sample buffer (2.0ml 1M Tris-HCl, pH 6.8, 2% SDS, 10% glycerol, 1% 14.7 M β-mercaptoethanol 12.5mM EDTA, 0.02% bromophenol blue). Proteins were then separated on 12% Bis-acrylamid mini gels and transferred onto PVDF membranes (Biorad, Sweden). The membranes were blocked for 60 min in 1% bovine serum albumin (in phosphate-buffered saline containing 1% Tween 20 (PBST); 137mM NaCl, 2.7mM KCl, 10mM Na_2_HPO_4_, 1.76mM K_2_HPO_4_; pH 7.4) and sequentially incubated with the primary and secondary antibodies. An antibody-specific dilution for the primary antibodies (**Supplementary Table 7**), and 1:40,000 for the HRP-labeled secondary antibody (BioNordika, Sweden) were utilized. All antibodies were diluted in 1% bovine serum albumin or 5% milk in PBS-T. A standard chemiluminescence procedure (ECL Plus) was used to visualize protein binding. The developed membranes were evaluated densitometrically using Image Lab 6.0.1 (Biorad, Sweden). All blots derive from the same experiment and were processed in parallel.

### Quantitative real-time PCR (qPCR)

RNA was isolated from brain tissue (N=5per group) using a Trizol® method. As per instructions using a “High-Capacity Reverse Transcriptase” kit (AB Bioscience, Stockholm, Sweden), 500ng of total RNA was reverse transcribed with random hexamer primers. The resulting cDNA was diluted to a final volume of 250μl and subsequently used as a template for qPCR reactions. qPCR was performed in triplicates using *Power* SYBR® Green PCR Master Mix (Life Technology, Sweden) according to the manufacturer’s instructions. Each reaction comprised 2ng cDNA, 3ul master mix and 0.2uM final concentration of each primer (see list of primers; **Supplementary Table 8**). Cycling and detection were carried out using a CFX Connect™ Real-Time PCR Detection System and data quantified using Sequence CFX Manager™ Software (Biorad, Sweden). qPCR was performed for a total of 40 cycles (95°C 15 sec, 60°C 60 sec) followed by a dissociation stage. All data were normalized to species specific housekeeping genes L14 (mouse) and GPI (human) and quantification was carried out via the absolute method using standard curves generated from pooled cDNA representative of each sample to be analyzed.

### Cell culture

Bone-marrow-derived macrophages (BMDMs) were generated by culturing freshly isolated mouse bone marrow cells from C57Bl/6J mice in IMDM medium (Gibco) supplemented with 10% FCS (GIBCO), 100 U/ml penicillin, and 100 U/ml streptomycin (Sigma-Aldrich), 2 mM glutamine and 20% (v/v) L929 conditional medium for 7 days. Human monocytic THP-1 cells (ATCC #TIB-202) were cultured in RPMI-1640 containing 10% fetal bovine serum, 0.05 mM β-mercaptoethanol, and 1 % penicillin-streptomycin in large culture flasks. Cell cultures were maintained at 37°C with 5 % CO_2_ and split 1:4 at a seeding density of 10^6^ cells. For differentiation, THP-1 cells were seeded in 6-well plates at a density of 3×10^5^ cells and treated with 2.5 ng/ml PMA for 48 hrs. Prior to LPS activation and treatment, cells were allowed to rest in culture media for 24 hrs.

Human peripheral blood mononuclear cells (PBMCs) were isolated using Ficoll-Paque Plus (GE healthcare) density gradient centrifugation from donated blood of healthy volunteers. After centrifugation, the PBMC layers were isolated and washed with PBS to remove erythrocytes and granulocytes. PBMCs purification was performed using anti-human CD14 magnetic beads (Miltenyi Biotec) according to the manufacturer’s instructions. Monocytes were maintained in RPMI 1640 (Gibco) with 10 % FCS (Gibco), 2 mM glutamine (Gibco), 100 U/mL penicillin and 100 µg/mL streptomycin (Gibco). For macrophage differentiation and maturation, human monocytes were plated onto tissue culture-treated 24-well plates at a density of 3×10^5^ cells/well. The medium was supplemented with 100 ng/mL recombinant human M-CSF using CellXVivo™ kit (R&D Systems) and differentiated for 5–7 days prior to use.

Cells were incubated with 1 µg/ml LPS (Invitrogen, Sweden) for 6 hrs prior to a 12 hrs treatment with 1 µM simvastatin (Bio-techne, UK), 10 µM hydralazine and/or a combination of both. The concentrations were chosen based on the dosing regimen administered in the *in vivo* studies. After incubation, cells were detached using cold PBS containing EDTA (5 mM) and either stained for flow cytometry or centrifuged and processed for RNA and protein isolation using the Trizol method.

### ELISA

TNF-α and IL6 protein levels in cell supernatant and plasma IL12/23 and IL17 protein levels were determined using DuoSet Elisa (R&D systems) as per manufacturer’s instruction.

### Statistical analysis

All data are expressed as mean ± SEM, where N is the number of animals or independent cell experiments. For comparisons of young and aged WT and ApoE^-/-^ mice, two-way ANOVA with Sidak post hoc testing was used to assess the effects of genotype and age. For comparison of multiple independent groups, the parametric one-way ANOVA test was used, followed by Tukey’s post hoc testing with exact P-value computation. In case of non-normally distributed data (tested with Shapiro-Wilk test), the non-parametric Kruskal Wallis test with Dunn’s post hoc testing and exact P-value computation was used for multiple comparisons. For comparison of two groups a two-tailed unpaired t-test was utilized. Differences were considered significant at error probabilities of P≤0.05. All statistical analyses were performed in GraphPad Version 8.1.2 (GraphPad Software, Inc).

## Data Availability

The data underlying this article are available in the article and in its online Supplementary Material.

## Acknowledgement

The authors thank Dr. Anna Planas for contributing aged ApoE^-/-^ mice and allowing us to use her behavioral platform, Francisca Ruiz for help with the treatment of the ApoE^-/-^ mice and the support with Elisa experiments, Dr. Harry Björkbacka for providing aged ApoE^-/-^ mice and access to his flow cytometer, Yun Zhang for help with running some of the flow cytometry, and Dr. Joao Duarte for sharing antibodies for SNAP-25 and PSD-95.

This work was supported by the following funding sources: The Knut and Alice Wallenberg foundation [2015.0030; AM]; Swedish Research Council [VR; 2017-01243; AM]; German Research Foundation [DFG; ME 4667/2-1; AM]; Åke Wibergs Stiftelse [M19-0380; AM]; Hedlund Stiftelse [M-2019-1101; AM], Inger Bendix Stiftelse [AM-2019-10; AM]; Stohnes Stiftelse [AM], Demensfonden [AM], Direktör Albert Påhlsson Stiftelse [AM], and startup funds provided by the Wallenberg Centre for Molecular and Translational Medicine, University of Gothenburg, Sweden [AH].

## Author Contributions

Conceptualization, AM; methodology, NDD, SR, LV, FM, HM and AM; validation, AM and AH; formal analysis, NDD, LV, SR and AM; data curation, NDD, SR, LV, FM, HM, SKP, AH and AM; writing—original draft preparation, AM; writing—review and editing, NDD, LV, HM, AH and AM; visualization, AM; supervision, AM and AH; project administration, AM; funding acquisition, AM and AH. All authors have read and agreed to the published version of the manuscript.

