## supplement for "Simvastatin Therapy Attenuates Memory Deficits that Associate with Brain Monocyte Infiltration in Chronic Hypercholesterolemia"

***SUPPLEMENTARY FIGURES***

***SUPPLEMENTARY FIGURE 1: Cholesterol- and BP-lowering therapy reduce systolic BP in aged ApoE$^{-/-}$ mice.*** Systolic BP of 12- and 14-months old ApoE$^{-/-}$ mice before and after treatment with BP- and cholesterol-lowering drugs (hydralazine, simvastatin and combination treatment). ApoE – apolipoprotein E, $BP_{sys}$ – systolic blood pressure. *Values expressed in mean ± SEM; N denotes number of independent biological replicates; N=10 for aged control, aged + hydralazine and aged + simvastatin treatment; N=8 for aged + combination treatment; * denotes P ≤ 0.05 relative 12 months (pre-treatment) after Two-way repeated measure ANOVA and Sidak post hoc testing.*

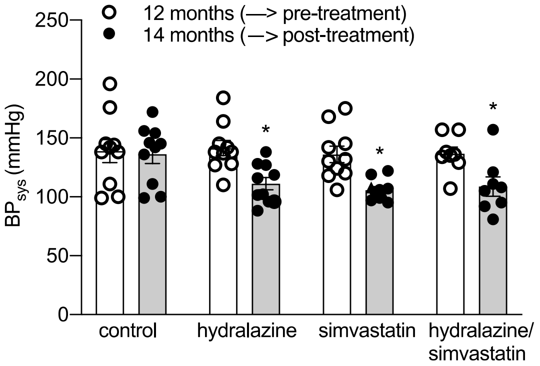


***SUPPLEMENTARY FIGURE 2: Region-specific microglia distribution and effect of treatment.*** **(A)** Total number of CD68+ Iba-1+ cells in the cortex and **(B)** total number of Iba1+ cells in the DG and **(C)** the cortex of 14-months old ApoE$^{-/-}$ mice compared to 4-months old ApoE$^{-/-}$ mice, and the effect of BP- and cholesterol-lowering therapy (hydralazine – green circle, simvastatin – pink circles, combination – orange circles). ApoE – apolipoprotein E, BP – blood pressure, DG – dentate gyrus, Iba-1 – ionized calcium-binding adapter molecule-1. *Values expressed in mean ± SEM; N denotes number of independent biological replicates; N=10 for aged + hydralazine; N=8 for aged control and aged + simvastatin treatment; N=6 for young control and aged + combination treatment; * denotes P ≤ 0.05 relative to young control, & denotes P ≤ 0.05 relative to aged control after Kruskal Wallis followed by Dunn's post hoc testing.*

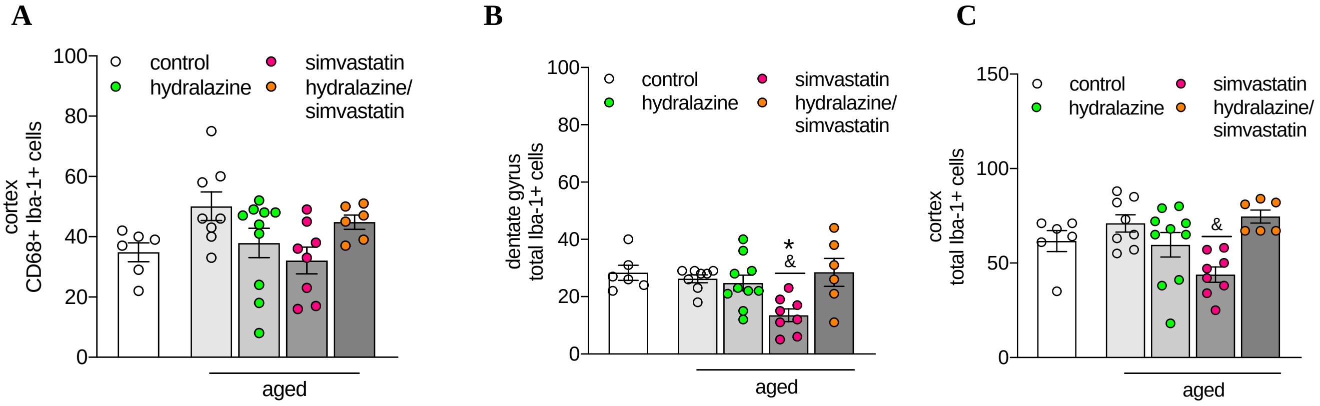

***SUPPLEMENTARY FIGURE 3: Simvastatin and Hydralazine treatment lower macrophage activation in primary mouse bone marrow-derived macrophages.*** Flow cytometry analysis of (**A**) CD86 and (**B**) CD80 responses to LPS, simvastatin and hydralazine in murine bone marrow-derived macrophages, and the effect of BP- and cholesterol-lowering therapy (hydralazine – green circle, simvastatin – pink circles). BP – blood pressure, LPS – lipopolysaccharide. *Values expressed in mean ± SEM; N denotes number of independent biological replicates, N=3 per group; * denotes P ≤ 0.05 relative to untreated control, & denotes P ≤ 0.05 relative to LPS control after one-way ANOVA followed by Tukey's post hoc testing.*

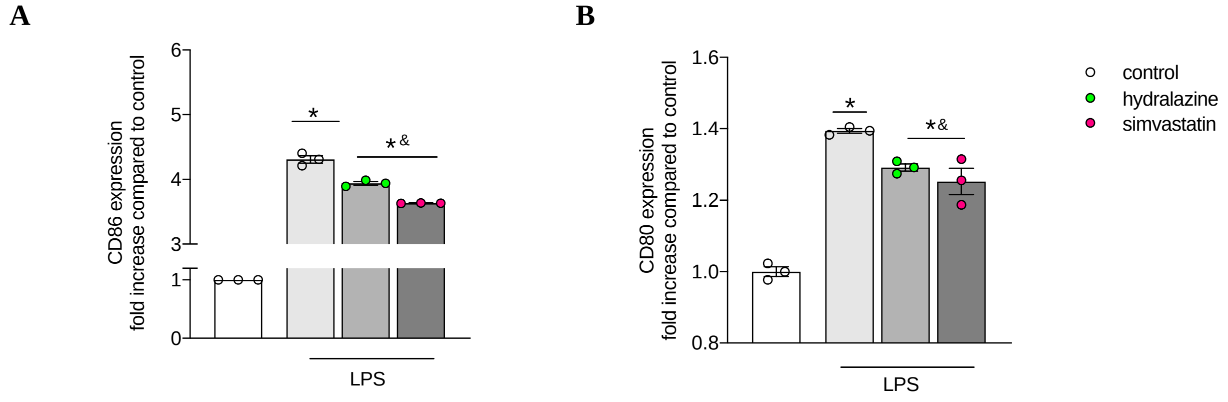


***SUPPLEMENTARY FIGURE 4: Hydralazine augments CD3 surface expression in human monocytic THP cells and primary human monocytes as well as monocyte-derived macrophages.*** Flow cytometric assessment of hydralazine effects on CD3 expression in (**A**) monocytic THP-1 cells, (**B**) PMA-differentiated THP-1 macrophages, (**C**) primary human blood monocytes, and (**D**) primary human monocyte-derived macrophages. **(E)** Representative histograms showing hydralazine responses in THP-1 cells and human primary cells. PMA - phorbol 12-myristate 13-acetate. *Values are expressed in mean ± SEM; N denotes number of independent biological replicates; N=3 per group; * denotes P ≤ 0.05 compared to respective control after unpaired t-test.*

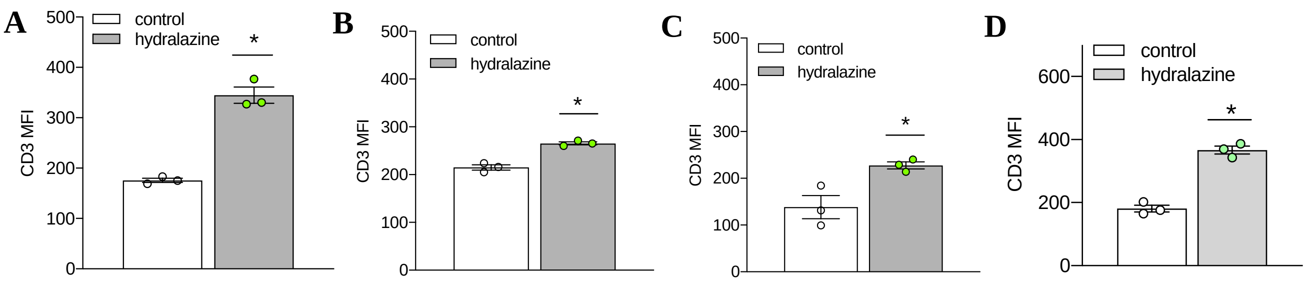

**E**

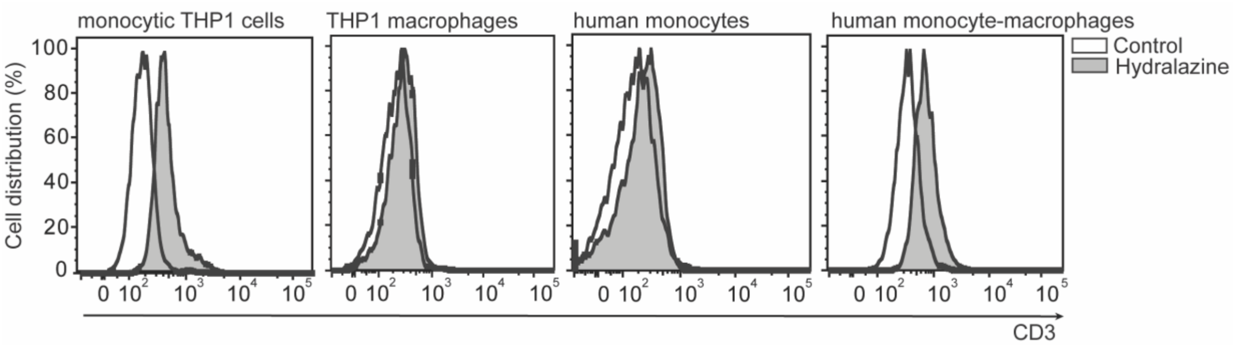


***SUPPLEMENTARY FIGURE 5: Simvastatin and Hydralazine treatment have no effect on Jurkat cell activation.*** Flow cytometry analysis of Jurkat cells showing percentage of CD69$^+$ cells following LPS, simvastatin or hydralazine treatment (hydralazine – green circle, simvastatin – pink circles, combination – orange circles). LPS –lipopolysaccharide. *Values expressed in mean ± SEM; N denotes number of independent biological replicates; N=4-7 per group; Significance tested with Kruskal-Wallis followed by Dunn's post hoc testing.*

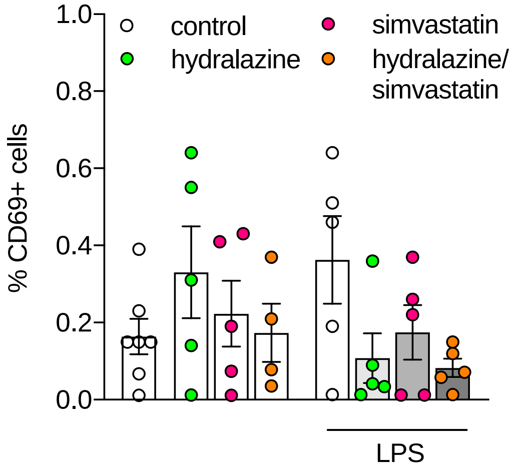


***<u>SUPPLEMENTARY TABLES</u>***

***SUPPLEMENTARY TABLE 1: Comparison of memory function in young and aged WT and ApoE$^{-/-}$ mice.*** Behavioral parameters of WT and ApoE$^{-/-}$ mice obtained in novel object recognition tasks with 5-min or 24-hrs delay interval. RI as the main index of retention were calculated by the time spent investigating the novel object relative to the total object investigation [RI = $T_{\mathrm{Novel}}/(T_{\mathrm{Novel}} + T_{\mathrm{Familiar}})$]. An RI=0.5 indicates that the animal has no preference for neither familiar nor novel object. ApoE – apolipoprotein E, RI – recognition index, WT –

wild-type. *Values expressed as mean+/- SEM; N denotes number of independent biological replicates; N=10 per group.* Mice with total object exploration times ($T_{Novel}$ + $T_{Familiar}$) below 20s were excluded from the analysis. *Statistical differences were assessed with non-parametric Mann Whitney tests for single unpaired comparisons.*

| | recognition index | P value | recognition index | P value |
|---|---|---|---|---|
| | 5 min delay | compared to young control | 24 hrs delay | compared to young control |
| WT young (4 months) | 0.71 ± 0.05 | | 0.57 ± 0.05 | |
| WT aged (12 months) | 0.74 ± 0.06 | 0.6403 | 0.58 ± 0.05 | 0.7974 |
| ApoE$^{-/-}$ young (4 months) | 0.76 ± 0.02 | | 0.63 ± 0.02 | |
| ApoE$^{-/-}$ aged (12 months) | 0.73 ± 0.03 | 0.4584 | 0.36 ± 0.04 | 0.0009 |

***SUPPLEMENTARY TABLE 2: Whole brain mRNA expression of inflammatory markers in 12-months old ApoE$^{-/-}$ versus 12-months old WT mice.*** ApoE – apolipoprotein E, CXCL – chemokine (C-X-C motif) ligand, GFAP – Glial fibrillary acidic protein, IL – interleukin, MCP-1 – monocyte chemoattractant protein 1, WT – wild-type. *Values are expressed as fold change compared to aged WT mice. N=5 per group; N denotes number of independent biological replicates.*

| gene | Fold change (ApoE$^{-/-}$ vs. WT) |
|---|---|
| MCP-1 | 5.673 |
| CXCL2 | 4.924 |
| GFAP | 1.969 |
| CD68 | 2.927 |
| IL6 | 5.507 |

***SUPPLEMENTARY TABLE 3: Effect of Simvastatin and Hydralazine treatment on NeuN and BDNF expression in different brain regions of ApoE$^{-/-}$ mice.*** Quantification of immune stained NeuN+ and BDNF+ cells in hippocampus regions (CA3, CA1 region and DG) and cortex from all experimental groups. ApoE – apolipoprotein E, BDNF – brain-derived neurotrophic factor, CA – cornu ammonis, DG – dentate gyrus *Values are expressed as mean +/- SEM; N=3 for young and aged + combination treatment; N=4 for aged control and simvastatin treatment; N=5 for hydralazine treatment. N denotes number of independent biological replicates. * denotes P ≤ 0.05 relative to young control, & denotes P ≤ 0.05 relative to aged group after one-way ANOVA followed by Tukey's post hoc testing.*

| | ApoE$^{-/-}$ young (4 months) | ApoE$^{-/-}$ aged (14 months) | ApoE$^{-/-}$ aged + hydralazine | ApoE$^{-/-}$ aged + simvastatin | ApoE$^{-/-}$ aged + hydralazine/simvastatin |
|---|---|---|---|---|---|
| hippocampus | | | | | |
| NeuN | 1078.2 ± 59.8$^{\&}$ | 558.5 ± 55.8* | 723.3 ± 69.8 | 797.5 ± 35.8*$^{\&}$ | 807.3 ± 50.1*$^{\&}$ |
| BDNF | 1484.0 ± 96.5$^{\&}$ | 438.5 ± 53.5* | 675.0 ± 73.9 | 935.6 ± 37.7*$^{\&}$ | 751.0 ± 67.3 |
| NeuN/BDNF ratio | 1.377 ± 0.054$^{\&}$ | 0.781 ± 0.027* | 0.953 ± 0.111* | 1.297 ± 0.125$^{\&}$ | 0.971 ± 0.041* |
| cortex | | | | | |
| NeuN | 768.0 ± 55.5 | 698.8 ± 55.9 | 940.8 ± 81.5 | 782.0 ± 39.3 | 939.0 ±50.2 |
| BDNF | 1650.0 ± 141.4 | 594.8 ± 68.0* | 1266.0 ± 105.2$^{\&}$ | 1139.0 ± 171.9$^{\&}$ | 1151.0 ± 182.8$^{\&}$ |
| NeuN/BDNF ratio | 2.142 ± 0.032 | 0.845 ± 0.035* | 1.364 ± 0.121 | 1.481 ± 0.265 | 1.213 ± 0.125 |

***SUPPLEMENTARY TABLE 4: Effect of hydralazine and simvastatin treatment on monocyte and macrophage activation.*** Flow cytometry analysis (percentages) of THP-1 cells, primary monocytes and monocyte-derived macrophages treated with hydralazine or simvastatin. *For each experimental group N=3. N denotes number of independent biological replicates. * denotes P ≤ 0.05 compared to control and & denotes P ≤ 0.05 compared to hydralazine group after one-way ANOVA followed by Tukey's post hoc testing.*

| | treatment | % CD3+ cells | % CD69+ cells |
|---|---|---|---|
| THP-1 monocytes | control | 1.08 ± 0.04 | 4.60 ± 0.40 |
| | 10uM hydralazine | 1.88 ± 0.09* | 6.90 ± 0.66* |
| | 1uM simvastatin | 1.16 ± 0.07$^{\&}$ | 3.33 ± 0.37$^{\&}$ |
| THP-1 macrophages | control | 1.03 ± 0.10 | 2.88 ± 0.25 |
| | 10uM hydralazine | 2.11 ± 0.14* | 4.53 ± 0.18* |
| | 1uM simvastatin | 1.27 ± 0.18$^{\&}$ | 2.40 ± 0.17$^{\&}$ |
| Human primary monocytes | control | 2.56 ± 0.23 | 1.70 ± 0.13 |
| | 10uM hydralazine | 4.50 ± 0.72* | 6.48 ± 0.38* |
| | 1uM simvastatin | 2.80 ± 0.35 | 2.30 ± 0.36 |
| Human primary monocyte-derived macrophages | control | 6.83 ± 0.33 | 8.07 ± 0.43 |
| | 10uM hydralazine | 16.00 ± 1.16* | 9.20 ± 0.51 |
| | 1uM simvastatin | 8.73 ± 0.62 | 6.33 ± 0.32* |

***SUPPLEMENTARY TABLE 5:*** **List of antibodies for flow cytometry.**

| Target | Host/species | Label/dye | clone | dilution | distributor |
|---|---|---|---|---|---|
| CD11b | Rat anti-mouse | PE-Texas Red | M1/70.15 | 1:100 | Life Technology |
| CD3 | Rat anti-mouse | eFluor450 | 17A2 | 1:100 | eBioscience |
| CD45 | Rat anti-mouse | AF700 | 30-F11 | 1:100 | eBioscience |
| CD45R | Rat anti-mouse | AF488 | RA3-6B2 | 1:100 | BioTechne |
| Ly6C | Rat anti-mouse | PE-Cy7 | HK1.4 | 1:100 | eBioscience |
| Ly6G | Rat anti-mouse | APC | RB6-8C5 | 1:100 | eBioscience |
| CD14 | Anti-human | eFluor450 | 61D3 | 1:100 | Life Technology |
| CD16 | Anti-human | FITC | CB16 | 1:100 | Life Technology |
| CD3 | Anti-human | FITC | BW264/56 | 1:100 | Miltenyi Biotec |
| CD69 | Anti-human | APC | 298614 | 1:100 | Life Technology |
| Live/dead | | Aqua-405 | | 1:250 | Life Technology |

***SUPPLEMENTARY TABLE 6:*** **List of antibodies used for immunofluorescence.**

| Antibody target | host | Supplier (catalogue #) | Antibody dilution |
|---|---|---|---|
| BDNF | rabbit | Abcam (ab6201) | 1:250 |
| NeuN | mouse | Abcam (ab104224) | 1:250 |
| Iba-1 | rabbit | WAKO (019-19741) | 1:1000 |
| CD68 (ED-1) | rat | Invitrogen (MA5-16654) | 1:500 |

***SUPPLEMENTARY TABLE 7:*** **List of antibodies used for Western blotting.**

| Antibody target | host | Supplier (catalogue #) | Antibody dilution |
|---|---|---|---|
| CD3 | rabbit | Abcam (ab5690) | 1:1000 |
| Beta-tubulin | mouse | Sigma Aldrich (T4026) | 1:3000 |
| SNAP-25 | rabbit | Abcam (ab109105) | 1:5000 |
| PSD-95 | rabbit | Abcam (ab76115) | 1:2000 |

***SUPPLEMENTARY TABLE 8:*** **List of mouse and human primers used for qPCR.**

| Target | Primer sequence (mouse) |
|---|---|
| L14 | F: 5'-GGCTTTAGTGGATGGACCCT-3’<br>R: 5'-ATTGATATCCGCCTTCTCCC-3' |
| TBP | F: 5’-GAAGCTGCGGTACAATTCCAG-3’<br>R: 5’-CCCCTTGTACCCTTCACCAAT-3’ |
| IL12 | F: 5'-CATCTGGTTCAGTGCTTTGATCT-3’<br>R: 5'-ACCCGTGAGTTATTCCATGAGT-3' |
| MCP-1 | F: 5'-CACGACATGCACGTTTGACT-3’<br>R: 5'-GGCGCAGATAAGGCTTCACA-3' |
| TNF-a | F: 5'-GTTTGCAGTCTCTCAAGCTTTT-3’<br>R: 5'-CCGATTTGAGCAATCGTTT-3' |
| IL6 | F: 5'-AGGAGTGATACCAGCTTTAGTCC-3’<br>R: 5'-CCGAGCAGGTCAGAACAAAGG-3' |
| iNOS | F: 5'-GGC AGC CTG TGA GAC CTT TG-3'<br>R: 5'-GCA TTG GAA GTG AAG CGT TTC-3' |
| CXCl2 | F: 5'-GTCCCAGACATCAGGGAGTAA-3’<br>R: 5'-TCGGATACTTCAGCGTCAGGA-3' |
| IL23 | F: 5'-GACCCACAAGGACTCAAGGAC-3’<br>R: 5'-ATGGGGCTATCAGGGAGTAGAG-3' |
| Arg-1 | F: 5'-TTGCGAGACGTAGACCCTGG-3’<br>R: 5'-CAAAGCTCAGGTGAATCGGC-3' |
| CD68 | F: 5'-TGTCTGATCTTGCTAGGACCG-3’<br>R: 5'-GAGAGTAACGGCCTTTTTGTGA-3' |
| CD86 | F: 5'-AGAACTTACGGAAGCACCCAC-3’<br>R: 5'-CTGCCAAAATACTACCAGCTCAC-3' |
| PSD-95 | F: 5'-ACCGCTACCAAGATGAAGACAC-3’<br>R: 5'-CTCCTCATACTCCATCTCCCC-3' |
| BDNF | F: 5'-TGCAGGGGCATAGACAAAAGG-3’<br>R: 5'-CTTATGAATCGCCAGCCAATTCTC-3' |
| **Target** | **Primer sequence (human)** |
| GPI | F: 5'-AGG CTG CTG CCA CAT AAG GT-3’<br>R: 5'-CCA AGG CTC CAA GCA TGA AT-3' |

| | |
|---|---|
| IFN-γ | F: 5'-ACTGACTTGAATGTCCAACGCA-3’<br>R: 5'-ATCTGACTCCTTTTTCGCTTCC-3' |
| IL-6 | F: 5'-TGC GTC CGT AGT TTC CTT CT-3’<br>R: 5'-GCC TCA GAC ATC TCC AGT CC-3' |

### Gating strategy

#### A. Brain

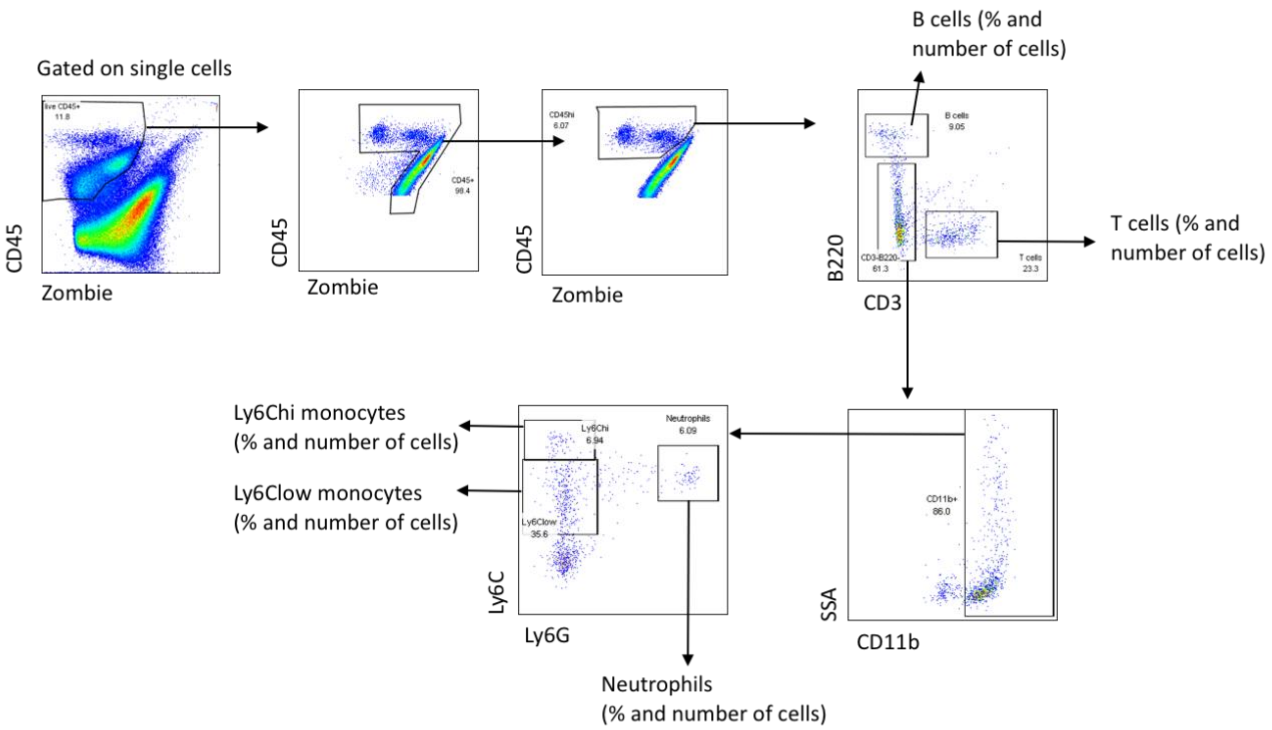


#### B. Blood

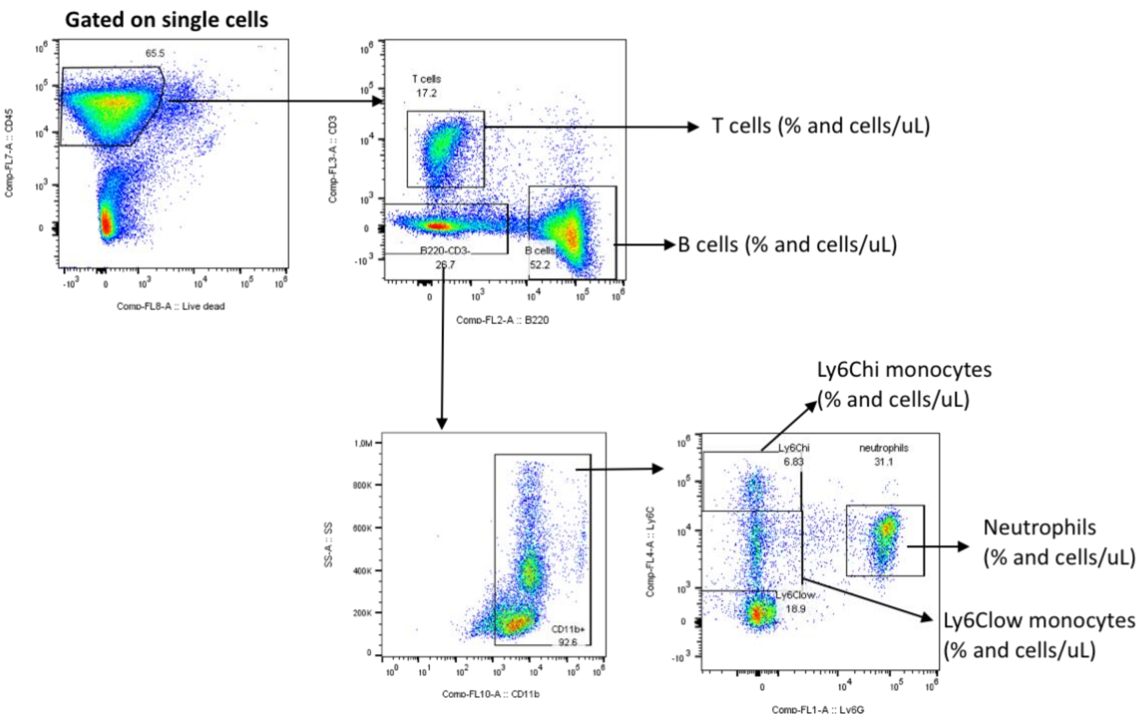

**Western blot images** (representative bands used for insets in main figures highlighted with red borders)

*Note: Full blots without cropping are given below. For SNAP-25 and PSD-95, molecular weight markers do not appear on the membranes because special stain-free markers were used in our system. To verify the protein sizes, a bright field image is obtained from the same membrane to overlay for analysis. The merged images are given below the membrane-only images. For CD3, the molecular weight marker is showing on the right corner of the membrane. The corresponding tubulin signal is too strong (1 sec exposure time) to visualize any protein standard band.

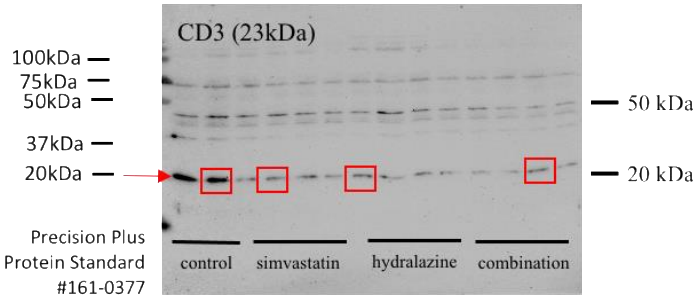


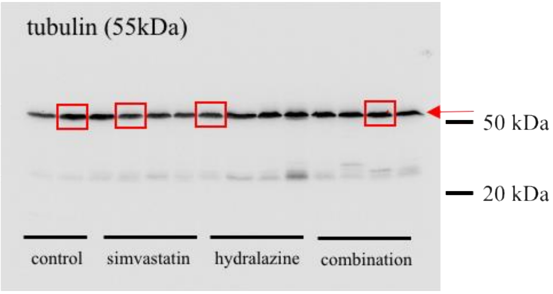


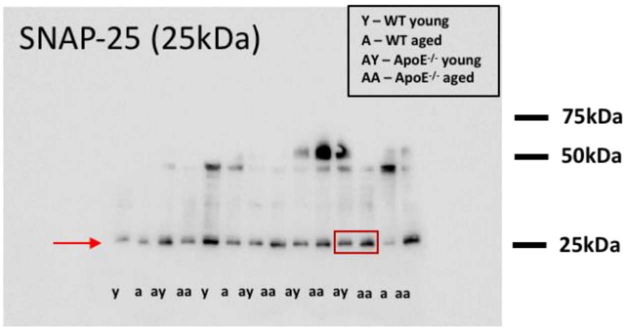


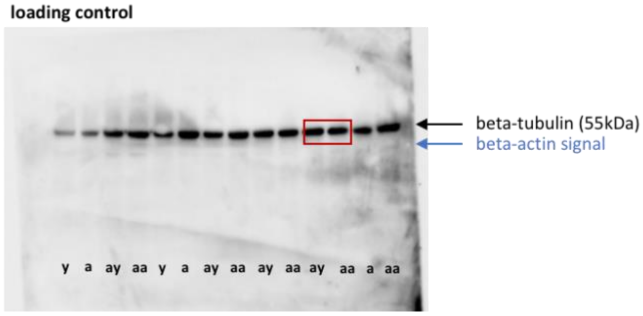


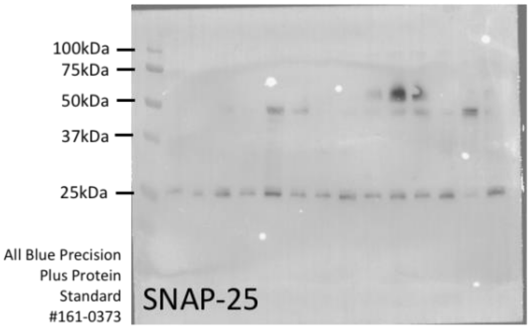


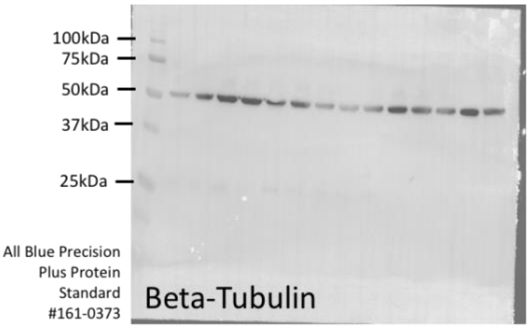

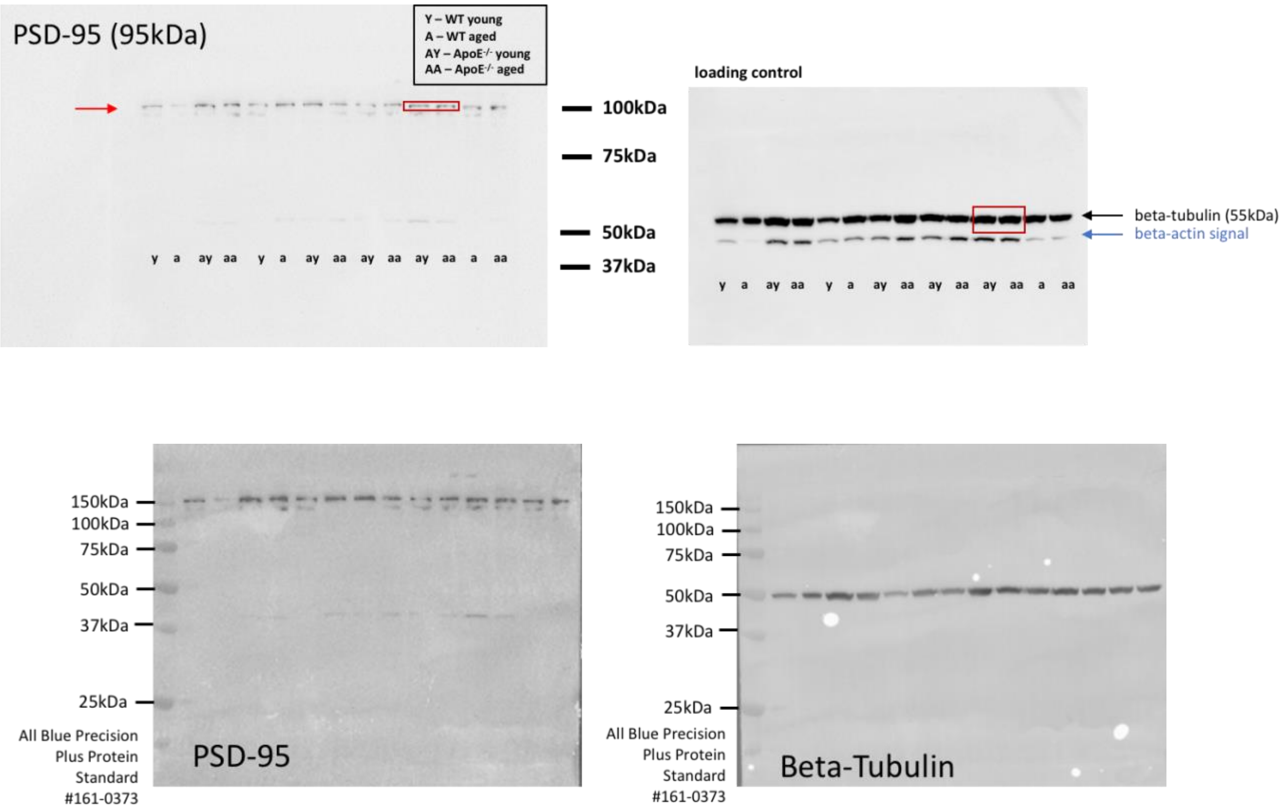
PSD-95 (95kDa)
Y – WT young
A – WT aged
AY – ApoE-/- young
AA – ApoE-/- aged
100kDa
75kDa
50kDa
37kDa
y a ay aa y a ay aa ay aa ay aa a aa
loading control
beta-tubulin (55kDa)
beta-actin signal
y a ay aa y a ay aa ay aa ay aa a aa
150kDa
100kDa
75kDa
50kDa
37kDa
25kDa
All Blue Precision Plus Protein Standard #161-0373
PSD-95
150kDa
100kDa
75kDa
50kDa
37kDa
25kDa
All Blue Precision Plus Protein Standard #161-0373
Beta-Tubulin
